# Cold induced chromatin compaction and nuclear retention of clock mRNAs resets the circadian rhythm

**DOI:** 10.1101/2020.06.05.127290

**Authors:** Harry Fischl, David McManus, Roel Oldenkamp, Lothar Schermelleh, Jane Mellor, Aarti Jagannath, Andre Furger

**Affiliations:** Department of Biochemistry, University of Oxford, South Parks Road, OX1 3QU; MRC LMB, Cambridge biomedical campus, Francis Crick Ave, Cambridge, CB2 0QH; The Netherlands Cancer Institute, Plesmanlaan 121, 1066 CX Amsterdam; Sir William Dunn School of Pathology, University of Oxford, South Parks Road, OX1 3RE

## Abstract

Cooling patients to sub-physiological temperatures is an integral part of modern medicine. We show that cold exposure induces temperature-specific changes to the higher-order chromatin and gene expression profiles of human cells. These changes are particularly dramatic at 18°C, a temperature synonymous with that experienced by patients undergoing controlled deep-hypothermia during surgery. Cells exposed to 18°C exhibit largely nuclear-restricted transcriptome changes. These include the nuclear accumulation of core circadian clock suppressor gene transcripts, most notably *REV-ERBα*. This response is accompanied by compaction of higher-order chromatin and hindrance of mRNPs from engaging nuclear pores. Rewarming reverses chromatin compaction and releases the transcripts into the cytoplasm, triggering a pulse of suppressor gene proteins that resets the circadian clock. We show that cold-induced upregulation of REV-ERBα alone is sufficient to trigger this resetting. Our findings uncover principles of the cellular cold-response that must be considered for current and future applications involving therapeutic deep-hypothermia.

## Introduction

Organisms are continually exposed to a variety of dynamic biological and environmental stressors. In order to survive, their cells must sense stresses and respond with structural alterations and the production of an adequate repertoire of metabolites and chaperones that enables them to reinstate cellular homeostasis and adapt to the new conditions (Kültz, 2020). Transcriptional reprogramming (Vihervaara *et al*, 2018) and post-transcriptional (Chua *et al*, 2020) adjustments of specific mRNA and protein levels in response to a particular stress is central to this adaptation. One of the most common abiotic stressors organisms experience result from changes in their environmental temperature. These temperature fluctuations have a profound impact on cells, affecting molecular structures, enzyme activity and the reactivity of metabolites, which can compromise viability.

We have a good understanding of how cells react to heat (Vihervaara *et al*, 2018; Al-Fageeh & Smales, 2006) but we know comparably very little about the gene networks and pathways that are activated when cells are exposed to cold (Ou *et al*, 2018). In bacteria, the gene expression response to cold stress is largely regulated at the post-transcriptional level where temperature-specific unfolding of mRNA structures by cold shock proteins (csps) regulates the global rate of translation initiation, elongation and mRNA stability. At the heart of this regulation is the chaperone activity of the cold shock protein cspA and its 5’UTR that acts as a thermosensor (Giuliodori *et al*, 2010) and auto-regulates its own abundance in a temperature-dependent way (Zhang *et al*, 2018). In plants, cold adaptation is more complex, involving temperature-induced adjustments of the chromatin (Zeng *et al*, 2019; Park *et al*, 2018), transcriptional (Nagano *et al*, 2019), pre-mRNA processing (Calixto *et al*, 2018), post-transcriptional and post-translational landscapes (Zhu, 2016). The transcriptional response to cold acclimatization in plants is beginning to be unraveled and entire signaling networks such as the C-repeat binding transcription factor (CBFs) regulon have been exposed (Zhao *et al*, 2015). Our understanding of how animal cells, especially mammalian cells, respond and adapt to cold temperatures is minimal and almost entirely limited to temperatures above 28°C (Al-Fageeh & Smales, 2006). Cold exposure to these temperatures activates the expression of the highly conserved cold-inducible RNA-binding protein (CIRBP) and RNA-binding motif protein 3 (RBM3) (Zhu *et al*, 2016). These two RBPs govern mammalian cold stress adaptations by regulating pre-mRNA processing and translation of gene transcripts that encode proteins with protective, including anti-apoptotic properties (Zhu *et al*, 2016). The protective effects attributed to RBM3 and CIRBP are of considerable medical interest (Peretti *et al*, 2015; Bastide *et al*, 2017) and exemplify the therapeutic potential that understanding of cold response mechanisms bear.

Despite considerable efforts to date, how cold responsive proteins are activated and how they ultimately confer cellular protection is largely unclear (Bastide *et al*, 2017). The regulatory networks are highly complex and have recently been shown to extend as far as the cellular circadian clock. The circadian oscillations in temperature that organisms experience are sufficient to activate both RBM3 and CIRBP, which in turn can modulate circadian gene expression post-transcriptionally by interacting with core clock gene transcripts (Morf *et al*, 2012; Liu *et al*, 2013). Cellular clocks enable cells to align their physiology with daily environmental cycles by controlling the rhythmic expression of thousands of genes. At the heart of the mammalian cellular clock lies a cell autonomous transcriptional feedback loop. The loop constitutes the core clock activators CLOCK and ARNTL (BMAL1) forming a heterodimeric complex that regulates the expression of E-box containing gene promoters, including those of the core clock suppressor genes: period 1 and 2, (PER1, PER2), cryptochrome 1 and 2 (CRY1, CRY2) and the nuclear receptor REV-ERBα (NR1D1) (Jagannath *et al*, 2017). Accumulation of these suppressor gene protein products inhibits the activity of the ARNTL/CLOCK complex, closing a feedback loop, resulting in the rhythmic expression of the core clock genes and oscillation of thousands of clock-controlled genes. This creates a complex interwoven regulatory network that enables individual cells and organisms to synchronize cellular physiology and organismal behavior with the daily solar cycle (Koike *et al*, 2012).

The activation of both RBM3 and CIRBP, however, is restricted to a very narrow temperature range between 28°C and 34°C (Rzechorzek *et al*, 2015). How mammalian cells respond to lower temperatures and how the activation of protective or detrimental pathways is controlled under such conditions is unknown (Ou *et al*, 2018). Work addressing cold responses in yeast (Kandror *et al*, 2004) and plants (Londo *et al*, 2018) suggest that different gene expression programs are activated depending on how low the temperature drops. Whilst most cells in warm-blooded animals are maintained within a narrow temperature range, cells and tissues of the extremities (Brajkovic & Ducharme, 2006) and cells of small hibernating mammals (Ruf & Geiser, 2015) regularly sustain temperatures below 10°C. Furthermore, human cells in internal organs are exposed to below 30°C during a number of medical procedures. These approaches exploit the reduction in cellular metabolic needs associated with cooling to minimize tissue damage resulting from ischemia when blood supply to vital organs is disrupted (Quinones *et al*, 2014). Current medical applications of controlled extreme cooling include exposure of organs or whole patients to as low as 18°C for a range of cardiothoracic and neurological surgeries conducted under deep hypothermic conditions (Mackensen *et al*, 2009). Transplant organs are routinely preserved by storage at temperatures below 10°C (Rao *et al*, 2001) and the body temperature of patients with major vascular damage are lowered to <10°C during pioneering emergency resuscitation procedures (Kutcher *et al*, 2016). Whilst the reduction of the metabolic needs of cells at very low (<28°C) temperatures are widely recognized (Ou *et al*, 2018), there remains a major conceptual gap in our understanding of the mechanisms and pathways that govern beneficial and destructive gene network adaptations to these conditions. Understanding these mechanisms will be critical to fully exploit the therapeutic potential of controlled therapeutic cooling (Kutcher *et al*, 2016).

Here we aimed to address the conceptual deficit of the mammalian cold stress response by tracking cold-induced changes to human cardiomyocytes exposed to temperatures (28°C, 18°C and 8°C) that are synonymous with the aforementioned medical procedures. We find that cold shapes both chromatin structures and subcellular transcriptomes in a temperature-specific manner. We discover that exposure to 18°C upregulates the transcript level of more than 1000 genes, including the core circadian clock suppressor gene transcripts *PER1, PER2, CRY1, CRY2* and *REV-ERBα*, but export of these transcripts into the cytoplasm is restricted. We provide evidence that the nuclear retention of the cold-induced transcripts at 18°C is caused by the compaction of chromatin into the nuclear periphery, compromising the approach of mRNPs to the nuclear pores. Export restrictions can be removed by rewarming cells back to 37°C, which reverses chromatin compaction and releases the accumulated nuclear transcripts into the cytoplasm. The sudden release of these mRNAs triggers a pulse of clock suppressor gene transcripts in the cytoplasm and their translation resets the circadian clock. Importantly, we show that REV-ERBα is the earliest to respond and most upregulated clock suppressor gene and its upregulation, following exposure to 18°C, is sufficient to reset the clock. Our findings expose a novel gene regulatory network and mechanism that links circadian gene expression with the mammalian response to very low temperatures.

## RESULTS

### Cold exposure to 18°C triggers temperature-specific, reversible changes to the higher-order chromatin architecture

To understand how mammalian cells respond to temperatures below the normo-physiological range (<30°C) that cells of internal organs experience during a number of medical procedures, we exposed human cardiomyocyte AC16 cells (Davidson *et al*, 2005) to 28°C, 18°C or 8°C. We then tracked temperature-dependent changes to the chromatin architecture by inspecting their 4’,6-diamidino-2-phenylindole (DAPI)-stained nuclei by super-resolution 3D-structured illumination microscopy (3D-SIM) (Schermelleh *et al*, 2008). This analysis revealed temperature-specific visible changes to the sponge-like fibrous structure formed by chromatin (Schermelleh *et al*, 2008) (Fig. 1A, S1A-C, DAPI panels). To quantify these topological changes, we applied a segmentation algorithm (Miron *et al*, 2019) that assigns each voxel of the nuclear DAPI signal into chromatin-depleted interchromatin (IC) and chromatin-enriched regions (Fig. 1A, segmented chromatin panels, Movies). As temperature does not affect the size of the nuclear volume (Fig. S1D), this reveals that at low temperatures chromatin undergoes compaction with a corresponding enlargement of the IC space. Surprisingly, this relationship is non-linear, as the greatest compaction and largest increase in IC occurs at 18°C, not 8°C (Fig. 1B, S1E). Upon rewarming back to 37°C, this cold-induced compaction to chromatin is reversed (Fig. 1B). Given the relationship between cold stress and chromatin modifications in plants (Kim *et al*, 2015), we next explored the possibility that the observed compaction may be accompanied by large-scale changes to the epigenetic landscape that may in turn regulate cold exposure-mediated cellular reprogramming. To assess this, we labelled the IC region as class 1 and further segmented the chromatin-enriched region into six classes from 2 to 7, denoting regions of increasing chromatin density. We then visualized the distribution of a number of chromatin features in cells exposed to the different cold temperatures across these seven chromatin density classes. These included serine 2-phosphorylated RNA polymerase II (Pol2S2P) and H3K4me3, features typically associated with actively transcribed chromatin, and H3K27me3 and H3K9me3 heterochromatin marks that can act as an impediment to cellular reprogramming (Nicetto & Zaret, 2019). After 24h at 28°C, 18°C or 8°C the heterochromatin marks remain enriched within the high-density chromatin classes 4-7 (Fig. S1A-B, S1F) and the active marks (Pol2S2P, H3K4me3) remain enriched within the low-density chromatin classes 2-3 (Fig. S1A-B, S1F). Thus, despite the global compaction of chromatin seen at low temperatures, including at 18°C, no cold-dependent large-scale changes in the epigenome topology or distribution of active transcriptional markers are evident in cold-exposed cells compared to cells kept at 37°C (Fig. S1F). This result suggests that the cold-induced chromatin compaction is mediated by physicochemical changes and is unlikely to be dependent on enzyme-driven modifications.

**Fig. 1.**
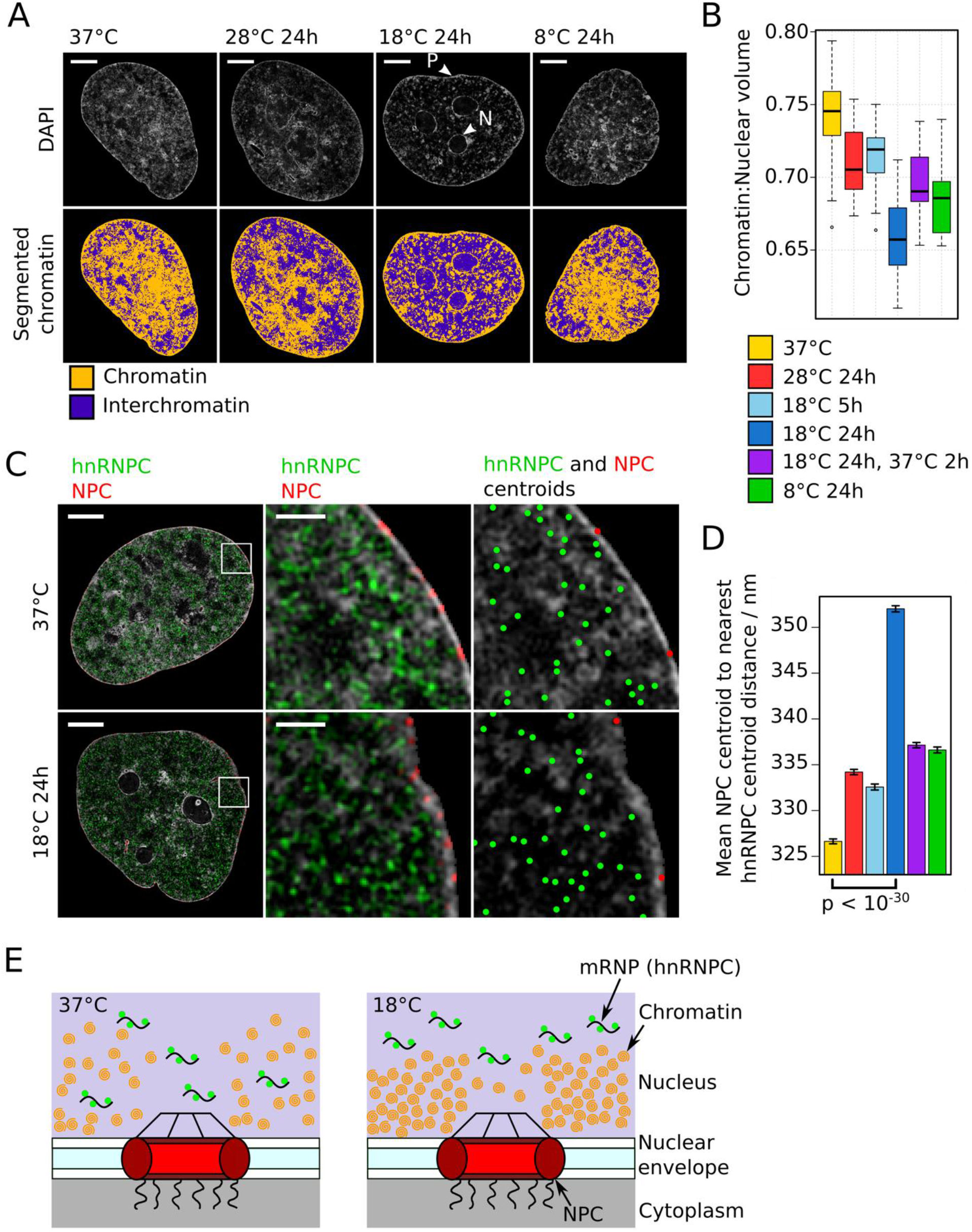
Cold temperature induces changes to the higher-order chromatin architecture. **A**) Upper panels: Representative single z-planes from 3D-SIM image stacks of the DAPI-stained nuclei of AC16 cells kept at 37°C or exposed to different cold temperatures for 24h. Lower panels: DAPI signal segmented into chromatin and interchromatin regions according to relative intensity. Scale bar: 10 μm. Arrows mark chromatin concentrated at the nuclear periphery (P) and around nucleoli (N). **B**) Boxplots of the ratio of chromatin volume to nuclear volume for cells exposed to different temperature conditions. **C**) Left and central panels: representative single z-planes from 3D-SIM image stacks of the DAPI-stained nuclei of cells kept at 37°C or exposed to 18°C for 24h showing the spatial distribution of the immunofluorescence (IF) signal of hnRNPC (green) and nuclear pore complexes (NPCs, red). Right panel: DAPI signal overlaid with dots indicating IF signal centroid coordinates for centroids located in the z-plane shown. Scale bar enlarged section: 1 μm. **D**) Bar graph of the mean NPC centroid to nearest hnRNPC centroid distance for each temperature condition. Error bars show the S.E.M. p-value (two-sided Welch’s t-test) calculated for the comparison shown. **E**) Schematic illustrating how chromatin concentrated at the periphery after exposure to 18°C could widen the average NPC to nearest mRNP distance and thus hinder mRNA nuclear export.

At 37°C, chromatin forms a diffuse network that is roughly evenly spread throughout the nucleus (Fig. 1A, S1A-C). In contrast, when chromatin compaction is at its greatest at 18°C, chromatin becomes visibly concentrated around nucleoli (“N”, Fig. 1A) and at the nuclear periphery (“P”, Fig. 1A), resulting in an apparent thickening of chromatin in these locations. To verify this thickening at the nuclear periphery, we visualized the location of nuclear pore complexes (NPCs) and hnRNPC molecules, which bind mRNAs in the IC space to form mRNA ribonucleoproteins (mRNPs) (Fig. 1C). As these two features localize to opposing sides of the lamin-associated chromatin at the nuclear periphery, changes in the average distance from the center of each NPC focus (centroid) to its nearest hnRNPC centroid are representative of changes in the thickness of this peripheral chromatin. Chromatin compaction at 18°C correlates with an increase in this average NPC to hnRNPC centroid distance (Fig. 1C-D, S1G-I). Upon rewarming, as chromatin at the periphery returns to a less compact state, there is a concomitant reduction in this average distance back towards levels seen in cells at 37°C (Fig. 1D). We conclude that 18°C cold exposure results in thickening of the chromatin at the nuclear periphery, which constrains the engagement of mRNPs with NPCs (Fig. 1E), and predict that this reduces mRNA export rates.

### Cold exposure shapes subcellular transcriptomes in a temperature-specific manner

We next addressed whether cold exposure at the aforementioned temperatures and the resulting compaction of chromatin shapes the nuclear and cytoplasmic transcriptomes. To that end, we used a subcellular fractionation approach (Neve *et al*, 2016) and sequenced the 3’ ends of nuclear and cytoplasmic polyadenylated RNAs from AC16 cells exposed to 28°C, 18°C or 8°C for 24h and compared their levels to those from cells kept at 37°C. Strikingly, each cold temperature results in its own specific effect on the subcellular transcriptomes (Fig. 2A). At 28°C, transcript level changes in the nucleus are generally mirrored by a change in the same direction and to a similar extent in the cytoplasm (Fig. 2B, top panel, S3D-E) with 651 and 1267 genes showing significant changes in the nucleus and the cytoplasm, respectively (Fig. 2A). In stark contrast, at 18°C, whilst over 2400 genes are significantly altered in the nucleus, cytoplasmic transcript levels are largely unaffected (Fig. 2A, B, middle panel). At 8°C, despite being the most extreme change in temperature, the nuclear transcriptome remains almost identical to that of cells kept at 37°C (Fig. 2B, bottom panel). Fewer than 70 genes show significant changes, suggesting transcription and nuclear RNA degradation and export are suspended. In contrast, cytoplasmic RNA levels do show changes at 8°C (Fig. 2B) with a significant reduction at more than 600 genes (Fig. 2A). Those showing greater decreases tend to have shorter half-lives (Lugowski *et al*, 2018) (Fig. S2A), suggesting cytoplasmic RNA degradation mechanisms are still active at 8°C. Transcriptome changes occurring in response to each cold temperature and within each compartment show only a weak positive correlation with those occurring in response to the other cold temperatures (Fig. 2C). Importantly, there is minimal overlap between the genes showing significant changes between the different temperatures (Fig. 2D). This is true for the known cold-inducible gene *CIRBP*, which is upregulated at 28°C but unaffected at 18°C and 8°C (Table S1, Fig. S2B). This confirms that it plays no part in the cellular adaptation process to very low temperatures. *RBM3*, which has also been shown to play a role in cold adaptation, is not upregulated at any of these cold temperatures (Fig. S2B). From these results, we conclude that the gene expression response to cold in mammalian cells is temperature and compartment-specific with little overlap between the gene networks that are activated at the different temperatures.

**Fig. 2.**
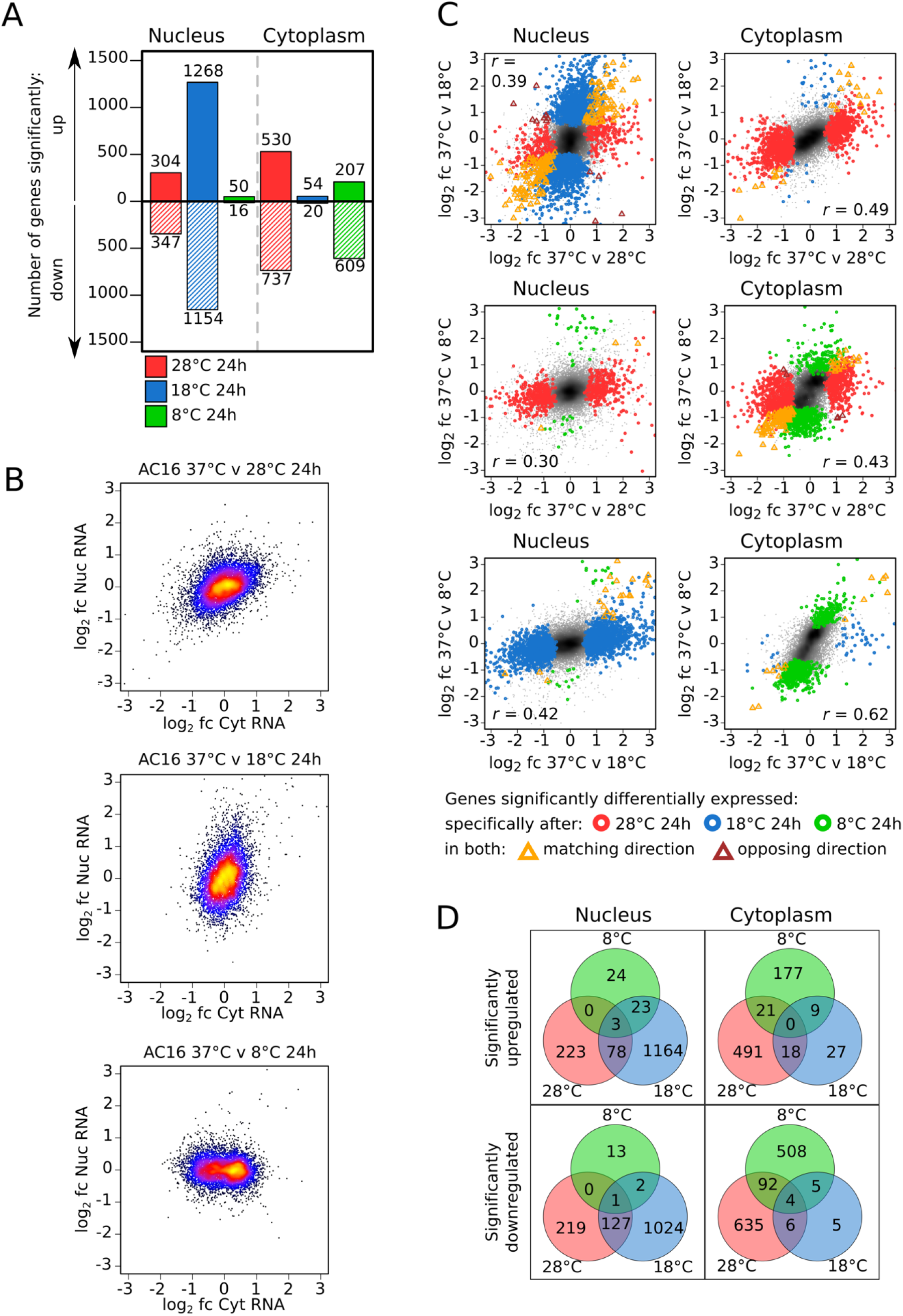
Cold induces temperature-specific changes to the nuclear and cytoplasmic transcriptome. **A**) Number of genes showing significant (adjusted p < 0.05) up or downregulation in nuclear or cytoplasmic RNA levels upon transfer of cells from 37°C to different cold temperatures for 24h. **B**) Relationship between the log2 fold change in RNA level in the nucleus and that in the cytoplasm for all genes upon transfer of cells from 37°C to different cold temperatures for 24h. Points are color-coded from low to high density (black < blue < red < yellow). **C**) Relationship between the log2 fold change in RNA level for all genes upon transfer of cells from 37°C to 28°C or 18°C (top), 28°C or 8°C (middle) and 18°C or 8°C (bottom) for 24h in both the nucleus and cytoplasm. Points are shaded from low to high density (grey < black). Genes that show significant differential expression at a specific temperature or at both temperatures are highlighted with colored circles or triangles, respectively. Pearson correlation coefficient (r_p_) is shown for each comparison. **D**) Venn diagram showing the overlap between the genes showing significant up or downregulation in nuclear or cytoplasmic RNA levels in cells exposed to the different cold temperatures.

### Upregulated transcripts at 18°C are nuclear-retained and are released into the cytoplasm by rewarming to 37°C

Emulating its impact on chromatin architecture, exposure to 18°C also has the most dramatic impact on the transcriptome. While over 1200 genes show upregulation of their nuclear RNA levels, their cytoplasmic levels remain unaffected (Fig. 2A-B). Importantly, spike-in controls confirm that these changes represent changes in absolute levels (Fig. S2C-E). The restriction of accumulated mRNAs to the nucleus suggests that whilst transcription is still active and responding to the 18°C cold exposure, nuclear-cytoplasmic mRNA export is compromised at this temperature. Furthermore, the disconnect between nuclear and cytoplasmic expression changes gradually intensifies and includes a growing number of genes the longer the cells are exposed to 18°C (Fig. S2F-H). Interestingly, rewarming cells back from 18°C to 37°C leads to a rapid readjustment of the nuclear-accumulated transcripts back to levels seen before cooling (Fig. 3A, top panel). This is concomitant with an increase of these transcripts in the cytoplasm (Fig. 3A, bottom panel). Likewise, the 1154 downregulated genes in the nucleus at 18°C are rapidly readjusted back to levels seen before cooling after rewarming (Fig. S3A). Moreover, these changes are not specific to AC16 cells, as the nuclear and cytoplasmic transcriptomes of U2OS cells, exposed to 18°C and then rewarmed, change in a similar way (Fig. S2E, S3B-C). This type of response is specific to 18°C as in cells that are exposed to 28°C, nuclear and cytoplasmic changes are mirrored during the entire cooling and rewarming cycle (Fig. S3D-E). Interestingly, the release of nuclear-accumulated transcripts into the cytoplasm after rewarming from 18°C coincides with the relaxing of the cold-induced compaction to chromatin (Fig. 1B), providing evidence that these events may be linked (Fig. 1E).

**Fig. 3.**
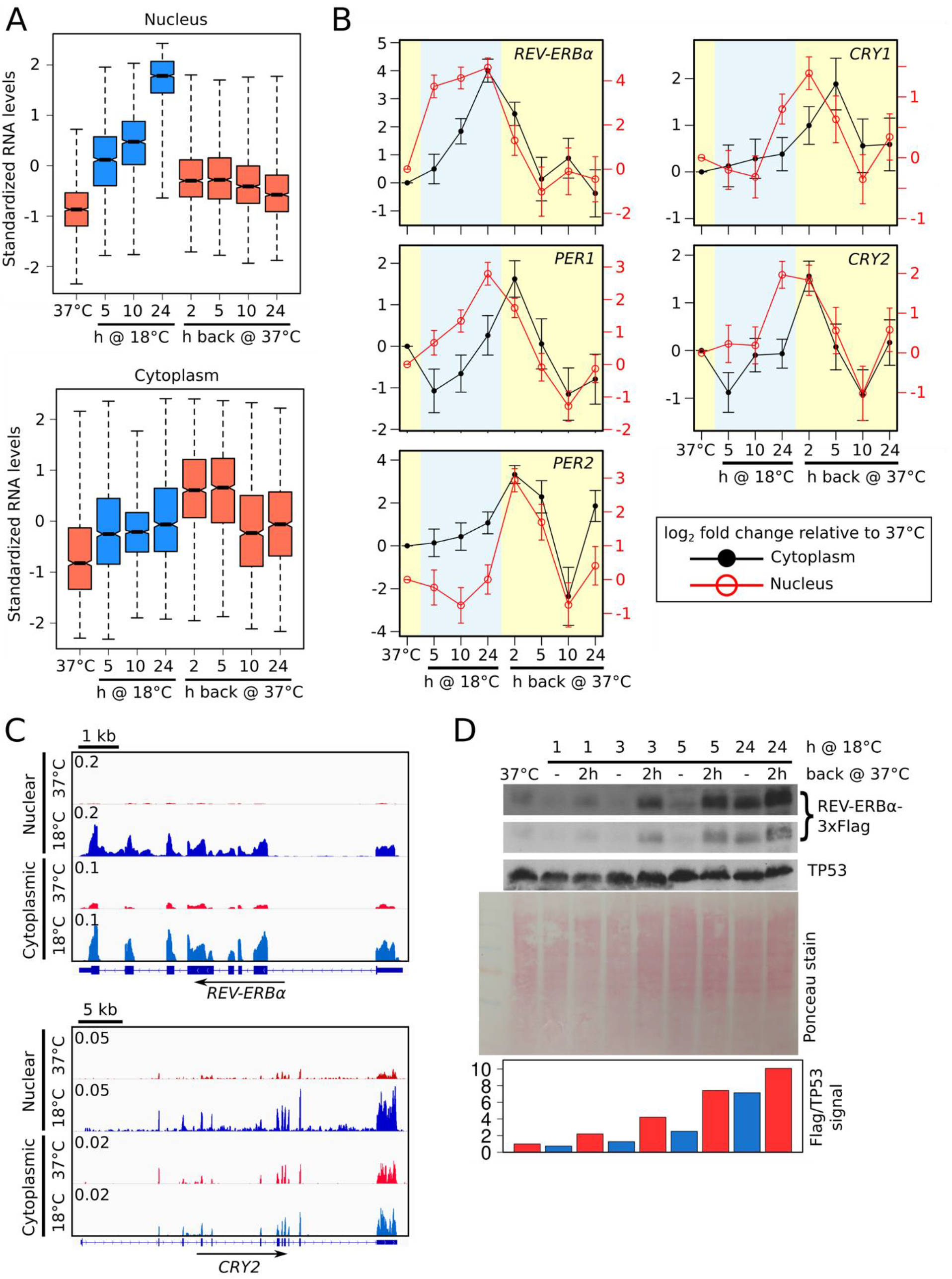
Rewarming following 18°C exposure releases nuclear-accumulated transcripts into the cytoplasm, including those of core circadian clock suppressor genes. **A**) Boxplots of standardized nuclear and cytoplasmic RNA levels at time points during the transfer of cells from 37°C to 18°C for 24h and then back to 37°C for 24h for the group of 1268 genes showing significant upregulation in nuclear RNA levels at the 18°C for 24h time point. **B**) Log2 fold change in cytoplasmic (black line, left axis) and nuclear (red line, right axis) RNA level of core circadian clock suppressor genes at time points during the transfer of AC16 cells from 37°C to 18°C for 24h and then back to 37°C for 24h relative to cells kept at 37°C. Error bars show the standard error. **C**) Nuclear and cytoplasmic full-length RNA-Seq data from AC16 cells kept at 37°C or exposed to 18°C for 24h for *REV-ERBα* and *CRY2*. Data for each track have been normalized to show counts per million aligned counts (CPM) for each nucleotide. Panels were prepared using Integrative Genomics Viewer (IGV). **D**) Western blot of Flag-tagged REV-ERBα levels (short and long exposure) in AC16 cells transferred from 37°C to 18°C for the time periods indicated and also returned after each of these time periods to 37°C for 2h. TP53 was used as a loading control as its transcript levels show minimal changes in response to 18°C exposure (Fig. 4, S3). Ponceau stain is shown as an additional loading control. Quantification of the REV-ERBα/TP53 signal is shown below.

### Cold exposure to 18°C triggers nuclear-restricted upregulation of core clock suppressor genes

We next explored the physiological consequence of this sudden release of nuclear-accumulated transcripts into the cytoplasm after rewarming cells from 18°C. To identify potential impacts, we subjected the cohort of transcripts that are upregulated in the cytoplasm after rewarming to gene ontology (GO) analysis, revealing an enrichment for genes associated with “circadian rhythm” (Table S2). Cellular circadian rhythmicity enables anticipatory alignment of physiological processes with daily environmental cycles with wide-ranging implications for health (Montaigne *et al*, 2018; Jagannath *et al*, 2017). As outlined earlier, these rhythms are orchestrated by a self-sustaining, cell-autonomous, molecular clock consisting of transcriptional activators, including ARNTL and CLOCK, and suppressors, including PER1/2, CRY1/2 and REV-ERBα, which interact in auto-regulatory feedback loops to generate oscillations in their expression levels that typically have 24h periodicity and distinct phases (Takahashi, 2017).

To explore the impact of the 18°C cold exposure on core clock gene expression in more detail we tracked the nuclear and cytoplasmic mRNA levels of these genes individually during exposure to 18°C and subsequent rewarming in AC16 (Fig. 3B, S3F) and U2OS cells (Fig. S3G). While the activators, *ARNTL* and *CLOCK*, do not show significant changes, suppressor components of the clock show nuclear-restricted accumulation of fully processed mRNAs (Fig. 3C). There are subtle differences in how these clock suppressor genes respond to 18°C. *REV-ERBα* accumulates rapidly in the nucleus after 5h exposure to 18°C and after 24h also seeps into the cytoplasm (Fig 3B). *CRY1, CRY2* and *PER1* increase after 24h at 18°C but the increase is nuclear-restricted (Fig. 3B). The nuclear-retained transcripts surge into the cytoplasm after rewarming to 37°C where they are translated (Fig. 3D, S3H). Unlike the other clock suppressor genes, *PER2* mRNA levels increase both in the nucleus and cytoplasm only after rewarming (Fig. 3B, S3G). This suggests that PER2 is activated by the rewarming process perhaps emulating a heat-shock response. However, it is unlikely that in our system rewarming induces *PER2* expression via the HSF1 pathway, as previously reported (Tamaru *et al*, 2011; Buhr *et al*, 2010; Kornmann *et al*, 2007), because we found no evidence that heat shock genes are activated during the cooling or subsequent rewarming process (Table S1).

To corroborate the observed RNA-Seq expression patterns of clock genes, we employed RNA-FISH using probes targeting transcripts for the most highly upregulated and earliest responding gene, *REV-ERBα*, the later responding and nuclear-restricted gene, *CRY2*, and the unchanged control gene, *TP53* (Fig. S3F). This analysis confirms that, after 5h at 18°C, *REV-ERBα* transcripts show rapid nuclear-restricted upregulation in both AC16 (Fig. 4A, E) and U2OS (Fig. 4D, H) cells, that, after 24h, *CRY2* transcripts accumulate in the nucleus and are released into the cytoplasm after rewarming (Fig. 4B, F), and that *TP53* transcript levels remain largely unchanged during cooling and rewarming (Fig. 4C, G).

**Fig. 4.**
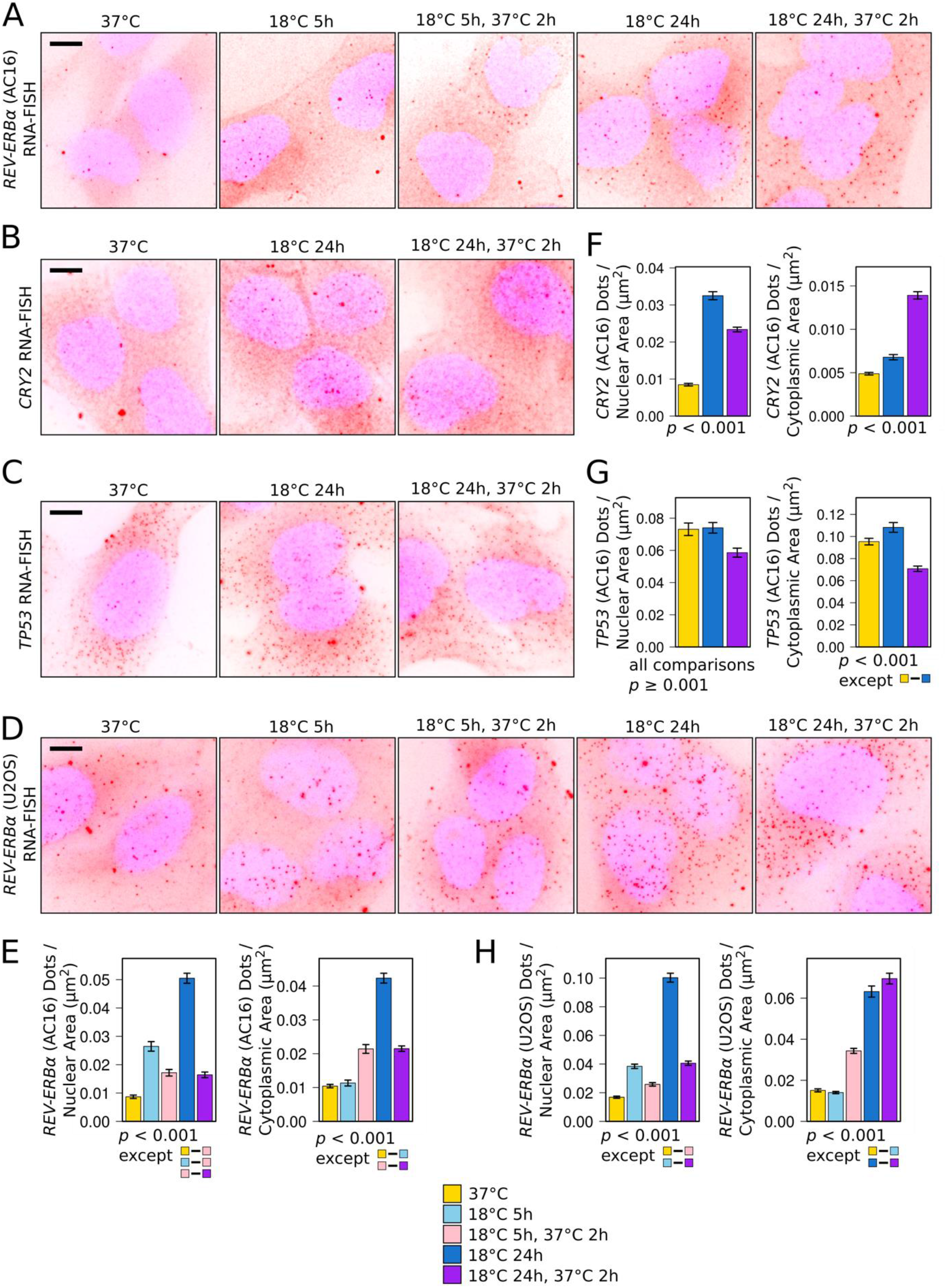
Nuclear accumulation upon 18°C exposure and release upon rewarming of core circadian clock suppressor gene transcripts. **A-D)** Representative maximum intensity z-stack projected images of DAPI-stained (blue) AC16 (A-C) or U2OS cells (D), exposed to 18°C and returned to 37°C for the time periods indicated, probed by RNA-FISH for *REV-ERBα* (A and D), *CRY2* (B) or *TP53* (C) transcripts (red dots). **E-H**) Mean *REV-ERBα* (E and H), *CRY2* (F) or *TP53* (G) transcripts per AC16 (E-G) or U2OS (H) cell nuclear and cytoplasmic area across all images for each condition from quantification of the RNA-FISH data. p < 0.001 (Tukey’s HSD) for all pairwise comparisons except those shown. All error bars (D-H) show S.E.M.

Collectively, these results show that the nuclear-restricted upregulation of the core clock suppressor gene transcripts is characteristic of the response to 18°C exposure and upon rewarming this nuclear pool of clock suppressor transcripts surges into the cytoplasm.

### Cold-induced upregulation of clock suppressor genes changes the phase and amplitude of the cellular circadian rhythm

We next investigated how this sudden burst of clock suppressor gene expression following rewarming affects the cellular circadian rhythm. In particular, we focused on the role of *REV-ERBα* given that, out of the clock genes, it is the only one to show upregulation after 5h, while also showing the greatest upregulation after 24h at 18°C (Fig. 3B). To monitor cellular circadian oscillations in real time, we tracked the bioluminescence generated by a *PER2* promoter-controlled luciferase reporter gene (*PER2::LUC*) stably expressed in U2OS cells with (WT cells) and without *REV-ERBα* (KO cells). In agreement with previous studies (Zhao *et al*, 2016), for cells kept at 37°C, *REV-ERBα* deletion increases the amplitude and period length of the circadian rhythm (Fig. S4A-B). We synchronized the cells with dexamethasone at regular intervals to produce cells with 6 distinct oscillating bioluminescence profiles distributed across the circadian period prior to cold exposure (6 differently colored lines, Fig. 5A, −48h-0h time window). This enables us to assess the relationship between the position of the cells within the circadian period at the point of cold exposure and any effect that the cold has on the subsequent amplitude and phase of the circadian rhythm. We first tested the effect of exposing cells to 18°C for 24h and subsequent rewarming, which results in elevated expression of *PER1/2, CRY1/2* and *REV-ERBα* (Fig. 3B). For all profiles, this causes a large significant increase in amplitude in both WT and KO cells (Fig. 5A, S4C, 24h-96h time window) compared to the amplitude of the profiles of cells kept at 37°C (Fig. S4G, S5C). In addition, it resets the clock by forcing cells into the same phase (clustered colored lines, Fig. 5A, S4C, 24h-96h time window) irrespective of the phase at which they oscillated before exposure to cold (Fig. 5A, S4C, G, S5B). To quantify this, for each profile we measured the time from the start of cold exposure to the second peak after rewarming and compared these to the time to the corresponding peak for profiles from cells kept at 37°C (peak “x”, Fig. 5C). The variance of these times for the different profiles is highly significantly reduced following exposure to 18°C for 24h, confirming that the phases have shifted, resulting in more closely aligned profiles (Fig. 5D). As this effect is independent of REV-ERBα levels (Fig. 5D, top panel: WT to KO cell comparison), we propose that the increased expression of the other clock suppressor genes is sufficient to reset the phase, masking any potential role for REV-ERBα.

**Fig. 5.**
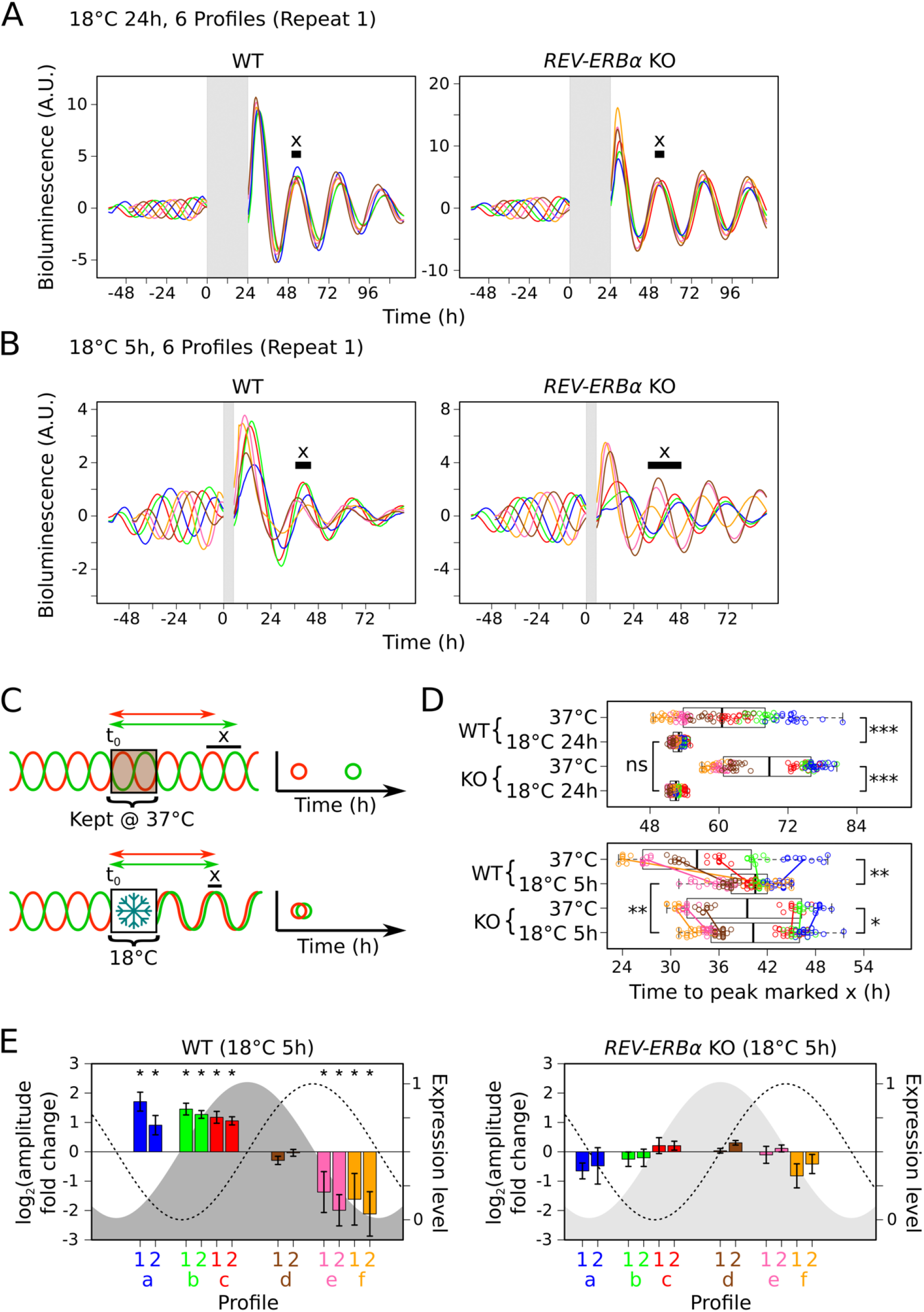
Cold-induced REV-ERB*α* expression resets the phase and modulates the amplitude of the circadian rhythm. **A** and **B**) Mean baseline-detrended bioluminescence profiles from plate wells containing either *PER2::LUC* U2OS cells with (WT, left panel) or without *REV-ERBα* (KO, right panel) recorded at 37°C before and after transfer to 18°C for 24h (A) or 5h (B) (grey region) from biological repeat 1. 6 differently colored profiles represent cells synchronized at 6 distinct phases of the circadian period prior to the start of 18°C exposure (time zero). **C**) Schematic showing the measurement of each profile from time zero to the peak marked “x” (this is the second peak after 18°C exposure or the corresponding peak in cells kept at 37°C (see A,B and Fig. S4C-E,G-I)). Plotting these times shows the effect of 18°C exposure on the spread of these profiles. **D**) Boxplots and individual points for times measured as in C for each plate well profile for both WT and KO cells kept at 37°C or transferred to 18°C for 24h (top panel) or 5h (bottom panel) from biological repeats 1 and 2. Points are colored according to their distinct phase prior to 18°C exposure. Colored lines (bottom panel) show the change in the mean for the points from each phase. Pairwise comparisons test the significance of the difference in variance (Brown-Forsythe test (adjusted for multiple testing)) (ns = p > 0.05; * = 1×10^−10^ < p < 0.05; ** = 1×10^−20^ < p < 1×10^−10^; *** = p < 1×10^−20^). **E**) Bar graphs showing the mean amplitude change following 18°C exposure for 5h for each group of plate well profiles (grouped according to their distinct phase (a-f) prior to 18°C exposure) for both WT and KO cells from biological repeats 1 and 2. Mean changes are plotted to approximately align with the position of the control profile at the point of rewarming within the circadian period that is represented by the *PER2:LUC* (dashed line) and predicted REV-ERBα expression level (grey shaded region (light grey indicates *REV-ERBα* deletion)). Error bars show the standard deviation. Asterisks mark mean log2 amplitude fold changes significantly greater than +/-log2(1.4) (adjusted p < 0.05). (see Fig. 5 extended data for more details).

Because rewarming following exposure to 18°C for 5h only triggers a cytoplasmic pulse of increased REV-ERBα without affecting *PER1/2* or *CRY1/2* levels (Fig. 3–4), we could use this to assess the effect of this cold-induced cytoplasmic surge of REV-ERBα on the circadian rhythm in isolation from the other clock suppressor genes. Upon rewarming following this shorter exposure, the phases of the WT cell profiles again become highly significantly more closely aligned than for cells kept at 37°C (Fig. 5B, left panel, Fig. 5D, bottom panel). Strikingly, this alignment now requires REV-ERBα expression as the profiles of KO cells do not align after cold exposure (Fig. 5B, right panel, Fig. 5D, bottom panel: WT to KO cell comparison; Fig. S4D, H; 4 additional profiles, Fig. S4E-F, I). To dissect the impact of elevated REV-ERBα levels in relation to its own rhythmicity, we calculated the phase and amplitude change for each profile (Fig. 5E, S5A, E-F) and plotted these in relation to the position of the cells in the circadian rhythm at the point of rewarming (Fig. S5D). This reveals that the REV-ERBα pulse forces a large significant phase delay when REV-ERBα (Fig. S5E, grey curve) should be decreasing (Fig. S5E: profiles d,y,e,f,z). This is greater for the profiles where REV-ERBα expression is nearing its minimum (Fig. S5E: profiles e,f,z), which also show large significant fold decreases in amplitude (Fig. 5E: profiles e,f; Fig. S5F: profile z). This may be because the minimum REV-ERBα level reached may no longer be as low due to the cold-induced pulse. Conversely, if the cold-induced pulse of REV-ERBα expression occurs when REV-ERBα should be increasing, there is a large significant fold increase in amplitude (Fig. 5E, profiles a,b,c; Fig. S5F: profiles w,x). This is likely because the maximum REV-ERBα level reached may now be higher. These results show that a single cold exposure to 18°C and subsequent rewarming creates a pulse in the expression of the core clock suppressor genes that changes the amplitude and resets the phase of the circadian rhythm. Importantly, cold-induced activation of REV-ERBα alone is sufficient to reset the phase.

Our results demonstrate that the core clock suppressor genes are part of a specific cellular response to very low temperature exposure that triggers resetting of the circadian rhythm upon rewarming.

## Discussion

Controlled cooling is an integral part of many clinical procedures, principally aiming to reduce the cellular metabolic needs of organs and tissues to prevent ischemic cell death when blood supply is interrupted. Whilst the benefit of reduced metabolic rates for cell survival has long been appreciated, the activation of specific pathways has only recently been recognized as a critical contributor to the cell preservation effect afforded by low temperatures (Bastide *et al*, 2017; Peretti *et al*, 2015; Ou *et al*, 2018) (Jackson & Kochanek, 2019). Unlike the metabolic rate that gradually reduces as temperature decreases, we show that the activation of pathways in response to cold is more complex. The mammalian response to cold is not scaled to temperature (Fig. 1–2) and is finely balanced between beneficial and detrimental outcomes (Hattori *et al*, 2017), limiting its therapeutic scope. The importance of controlled cooling as a clinical tool (Peretti *et al*, 2015; Gordon, 2001; Bastide *et al*, 2017; Kutcher *et al*, 2016) and its potential to provide solutions to overcome the physiological challenges of long duration space flight (Nordeen & Martin, 2019; Choukèr *et al*, 2019; Jackson & Kochanek, 2019), expose the need to understand the cellular response to cooling at the molecular level. Elucidating these processes will ultimately provide solutions to pharmacologically activate beneficial pathways without the need for cooling and inactivate detrimental pathways to expand the remit of controlled cooling for medical applications.

Here we describe a multipronged approach to elucidate how cells respond to cold temperatures that are currently applied in a number of clinical settings. Using this approach, we find that, similarly to plants and yeast, human cells respond to cold in a highly temperature-dependent manner and that the activation of core clock genes is an integral part of the mammalian response to extreme cold temperature exposure.

The temperature-dependent cellular response is most apparent in the dramatic compaction of chromatin in cells exposed to cold, particularly at 18°C (Fig. 1). Surprisingly, our 3D-SIM approach shows that the degree of compaction is non-linear and independent of enzymatic activity and is thus most likely caused by a physicochemical change in the nucleus. A probable trigger for this change is a temperature-induced influx of calcium that is stored in the nuclear envelope and the nuclear reticulum (Bootman *et al*, 2009). Not only is calcium known to boost compaction of chromatin fibers (Phengchat *et al*, 2016) but calcium release channels are activated in a temperature-dependent manner, operating as thermo-sensors in both mammalian (Bautista *et al*, 2007) and plant cells (Ding *et al*, 2019).

The use of the nuclear cytoplasmic fractionation approach added a novel dimension to the mammalian cold response by identifying the establishment of distinct temperature-specific nuclear and cytoplasmic transcriptomes (Fig. 2). These differences are due to temperature-specific effects on mechanisms controlling transcription, mRNA export and mRNA degradation. The activation of specific programs depending on how low the temperature drops results in gene expression outputs with very little overlap between the affected genes at the different temperatures (Fig. 2C-D). At 28°C, changes compared to cells kept at 37°C are characterized by the upregulation of around 500 genes, including the known cold-induced gene *CIRBP*. The changes in the nuclear and cytoplasmic transcriptomes are generally matched, suggesting that the predominant response at 28°C is at the transcriptional level. This is in marked contrast to 8°C where the nuclear transcriptome is indistinguishable from that of cells kept at 37°C. This may be caused by a depletion of ATP at this temperature which will impair transcriptional rates (Pires *et al*, 2010), reduce the free movement of mRNPs away from chromatin and prevent their nuclear export (Vargas *et al*, 2005), essentially “freezing” the status quo in the nucleus. In contrast, cytoplasmic degradation pathways are still active leading to greater depletion of unstable transcripts (Fig. S2), causing the disparity between nuclear and cytoplasmic 8°C transcriptomes (Fig. 2).

The most dramatic transcriptome change is associated with shifting cells to 18°C, which causes a striking upregulation and retention of over 1000 genes within the nucleus (Fig. 2–4). Incomplete pre-mRNA processing has been identified as a key determinant for nuclear transcript retention (Yin *et al*, 2020) but this mechanism is unlikely to be responsible for the cold-induced restriction to export because the upregulated retained transcripts are polyadenylated and no evidence for increased intron retention was observed (Fig. 3C). Furthermore, the cytoplasmic surge of the upregulated transcripts into the cytoplasm and their translation (Fig. 3D) after rewarming confirms that these transcripts are fully matured. Our 3D-SIM and RNA-FISH data (Fig. 1) suggest that the nuclear accumulation is linked to the high degree of chromatin compaction at 18°C. We propose that the cold-induced thickening of peripheral chromatin hinders mRNPs from approaching the nuclear pores along the chromatin free channels (Schermelleh *et al*, 2008) and that the compaction of the lamin-associated chromatin together with the low temperature affects the rigidity of the membrane, altering the mechanical properties of the NPCs. Collectively these effects result in almost complete inhibition of nuclear cytoplasmic mRNA export, creating the highly polarized subcellular transcriptomes characteristic for this temperature (Fig. 1E). Importantly, these effects are entirely reversible and rewarming prompts relaxation of the chromatin and release of the accumulated transcripts into the cytoplasm.

Interestingly, rewarming cells from 18°C rapidly reverts the chromatin configurations and readjusts both the nuclear and cytoplasmic transcriptomes to precooling states (Fig. 3–4). With the exception of a few genes, this process takes less than 2h for the nuclear transcriptome and less than 10h for the cytoplasmic transcriptome (Fig. 3A), suggesting a high degree of coordination between mechanisms that control the rates of transcription and degradation during this short period.

A key finding of our detailed transcriptome analysis is the recognition that the core clock suppressor genes, *PER1, CRY1/2* and *REV-ERBα* can be classified as cold response genes. The nuclear-restricted upregulation of these genes is dramatic, most notably for *REV-ERBα*, which exceeds a greater than 10-fold change in transcript abundance at 18°C compared to levels in cells kept at 37°C (Fig. 3B). Importantly our data demonstrates that the upregulation and the cytoplasmic burst of the clock suppressor genes into the cytoplasm following rewarming, forces a phase and amplitude change in the circadian rhythm (Fig. 5). This is a critical finding as it demonstrates the observed transcriptome changes are of physiological importance and uncovers a novel mechanism by which temperature influences the cellular circadian clock. There is an intricate relationship between the circadian clock and temperature. For example, repeated cycling between high and low temperatures within the narrow physiological temperature range typically experienced by homeothermic mammals (33-37°C) is sufficient to entrain circadian rhythms (Brown *et al*, 2002). RBM3 and CIRBP, activated by such mild temperature changes, are key regulators of this process (Liu *et al*, 2013; Morf *et al*, 2012). Similarly, exposing cells to a heat shock causes circadian rhythms to reset in a HSF1-dependent manner (Buhr *et al*, 2010; Tamaru *et al*, 2011). However, both of these pathways cannot account for the observed phase changes associated with exposure to 18°C and subsequent rewarming as neither *CIRBP, RBM3* nor *HSF1* are upregulated (Table S1). Instead, we identified a novel mechanism by which a phase change is triggered by a cold-induced impairment of nuclear export of upregulated clock transcripts and their sudden release into the cytoplasm after rewarming. How the expression of the clock suppressor genes is upregulated in the nucleus is unclear but may be dependent on an influx of calcium that could activate calmodulin–CaMKII kinase and so boost E-box gene expression by promoting dimerization of CLOCK and ARNTL (Kon *et al*, 2014).

Our data further show that the cold-induced upregulation of REV-ERBα alone is sufficient to trigger a phase shift. This is of high significance as it not only identifies REV-ERBα as an important cold-induced gene but it also demonstrates that massive upregulation of REV-ERBα is sufficient to control phase resetting in mammalian cells. Finally, given that REV-ERBα is a key chromatin regulator (Kim *et al*, 2018) linking circadian rhythms with metabolism (Cho *et al*, 2012; Zhang *et al*, 2015), inflammation (Pariollaud *et al*, 2018) and cardiac surgery outcomes (Montaigne *et al*, 2018), and has tumor suppressor (Sulli *et al*, 2018) and anti-viral properties (Zhuang *et al*, 2019), the recognition that *REV-ERBα* expression can be boosted more than ten-fold by 18°C exposure will be of significance for the development of novel strategies that broaden the applications of controlled deep hypothermia.

## Material and Methods

### Cell culture

AC16 (Davidson *et al*, 2005) cells were cultured in Dulbecco’s Modified Eagle’s Medium: Nutrient Mixture F-12 (DMEM/F-12), containing 10 % (v/v) Foetal Bovine Serum (FBS), 2.5 mM L-Glutamine and 1 % Penicillin/Streptomycin (Sigma cat#P0781).

We used a U2OS osteosarcoma cell line stably expressing the luciferase reporter gene (*luc2* (*Photinus pyralis*)) under the control of the mouse *PER2* promoter (U2OS *PER2::LUC*) (Jagannath *et al*, 2013). U2OS cells were cultured in DMEM containing 10 % (v/v) FBS, 2.5 mM L-Glutamine and 1 % Penicillin/Streptomycin (Sigma cat#P0781) except during bioluminescence recording (see below).

Unless otherwise specified, human cells were cultured in a 37°C incubator with 5 % CO2 to 80-90 % confluency before harvesting, passaging or transfer to incubators set at 8°C, 18°C or 28°C (also with 5 % CO2).

*D. melanogaster* Schneider 2 (S2) cells were cultured in Schneider’s Drosophila Medium (ThermoFisher, cat.no. 21720024), 10% heat-inactivated FCS and 1 % Penicillin/Streptomycin (Sigma cat#P0781) in a 28°C incubator with 5 % CO2 to 80-90 % confluency before harvesting. Cells were aliquoted, pelleted and flash frozen. Aliquots harvested at the same time were used in all spike-in experiments.

### Genome editing

#### *REV-ERBα* deletion

sgRNAs were cloned into the pSpCas9(BB)-2A-Puro(PX459)-V2.0 vector (Addgene #62988) by annealing oligonucleotides 1 and 2 (Table S3) and inserting them into the BbsI site, as previously described (Ann Ran *et al*, 2013). For *REV-ERBα* knock out, two sgRNAs, sgRNA Del US and sgRNA Del DS, were designed to cleave 723 bp upstream and 135 bp downstream of the start codon, respectively, to create an approximate 878 bp deletion of a region containing the promoter and first exon of *REV-ERBα*. This leads to a frame shift of subsequent exons, disrupting REV-ERBα function. Cloned vectors (1.5 μg each per 6 well plate well) were transfected using Lipofectamine 3000 (ThermoFisher) according to the manufacturer’s instructions into U2OS *PER2::LUC* cells. As this cell line is already puromycin resistant, no selection for transfected cells was possible. Individual colonies were screened by PCR to identify correctly targeted homozygous clones.

#### *REV-ERBα*-3xFLAG tagging

For *REV-ERBα* 3xFLAG C-terminal tagging, one sgRNA, sgRNA Tag was designed to cleave 3 bp downstream of the stop codon. The cloned vector (3.5 μg) was transfected into AC16 cells with 1.5 μl single-stranded oligonucleotide (ssODN) (10 μM) as a template for homology-directed repair (HDR)-mediated 3xFLAG tag knock-in (Table S3). The ssODN was designed with asymmetric homology arms, which have been proposed to enhance HDR efficiency (Richardson *et al*, 2016). Transfected cells were selected by incubation with puromycin (1 μg/ml) in DMEM for 48 h and individual colonies were screened by PCR and Western blot to identify correctly targeted homozygous clones.

### Nuclear and cytoplasmic RNA extraction

Extraction of RNA from nuclear and cytoplasmic subcellular fractions was carried out as described in (Neve *et al*, 2016). 1 x 10^7^ AC16 cells were trypsinized and washed twice in ice cold PBS. Cytoplasmic membranes were lysed by slowly resuspending the cell pellet in 1 ml Lysis Buffer A (10 mM Tris-HCl (pH 8-8.4), 0.14 M NaCl, 1.5 mM MgCl2, 0.5 % NP-40). Nuclei were pelleted (1000 g, 4°C, 3 min) and the supernatant, consisting of the cytoplasmic fraction, was cleared of remaining nuclei by spinning (11000 g, 4°C, 1 min). The pelleted nuclei were resuspended in 1 ml Lysis Buffer A. 100 μl Detergent Stock Solution (3.3 % (w/v) sodium deoxycholate, 6.6 % (v/v) Tween 40) was then added dropwise under slow vortexing. Stripped nuclei were pelleted (1000 g, 4°C, 3 min), washed once with 1 ml Lysis Buffer A, and then resuspended in 1 ml Trizol (ThermoFisher). 500 μl of Trizol was also added to 500 μl of the cleared cytoplasmic fraction. RNA was purified from the nuclear and cytoplasmic fractions as per the Trizol manufacturer’s instructions. RNA samples were DNase I-treated (Roche) as per the manufacturer’s instructions, purified using phenol-chloroform and precipitated with ethanol.

For *D. melanogaster* Schneider 2 (S2) cell spike-in experiments, S2 and AC16 cells were mixed at a known ratio (either 1:4 (replicates 1 and 2) or 5:4 (replicates 3 and 4)) before cell lysis.

### RNA-Seq

The number of replicates sequenced for each condition are listed in Table S4.

#### QuantSeq

Barcoded libraries for multiplexed, strand-specific sequencing of the 3’ end of polyadenylated RNAs were generated using the QuantSeq 3’ mRNA-Seq Library Prep Kit for Ion Torrent (Lexogen) as per the manufacturer’s instructions, using 500 ng and 1700 ng input RNA for nuclear and cytoplasmic RNA samples, respectively, and using 13 PCR cycles. Libraries were loaded onto the Ion Chef System (ThermoFisher) for template preparation and chip loading and the resulting chips were sequenced on the Ion Proton Sequencing System (ThermoFisher) as per the manufacturer’s instructions.

For standard libraries, reads were aligned to the hg19 genome build using the Ion Torrent Server TMAP aligner with default alignment settings (-tmap mapall stage1 map4). Human polyA site (PAS) annotations were obtained from PolyA_DB 3 (http://exon.umdnj.edu/polya_db/v3) (Wang *et al*, 2018). Each PAS was extended 20 nt 3’ and 200 nt 5’ from the site of cleavage and those that overlapped on the same strand after extension were combined into a single PAS annotation. Mapped reads were narrowed to their 3’ most nucleotide and those which overlapped with the extended PAS annotations were counted. Counts were then obtained for each gene by combining the counts for all PASs associated with each gene. Genes not in the RefSeq gene database were excluded.

For libraries prepared from *D. melanogaster* Schneider 2 (S2) cell spiked-in samples, reads were simultaneously aligned to both human and *D. melanogaster* genomes by aligning to a custom combined hg19 and dm6 genome build using the TMAP aligner. *D. melanogaster* PASs were obtained from the Tian lab (Liu *et al*, 2017) and were extended, in the same way as for human PASs. Mapped reads were narrowed to their 3’ most nucleotide and those which overlapped with the extended human or *D. melanogaster* PAS annotations were counted. The ratio of total *D. melanogaster* PAS to human PAS associated reads was obtained for each sample. These were then scaled relative to each other according to the ratio at which the two cell lines were combined.

#### Full-length RNA-Seq

Barcoded libraries for multiplexed, full-length, strand-specific sequencing of polyadenylated RNAs were generated using the Ion Total RNA-Seq Kit v2 (ThermoFisher) as per the manufacturer’s instructions. Polyadenylated RNAs were first purified from total RNA from each fraction using the NEBNext Poly(A) mRNA Isolation Module as per the manufacturer’s instructions. Libraries were sequenced and aligned in the same way as the QuantSeq libraries. Data were normalized to sequencing depth by calculating the count at each nucleotide per million counts aligned at all nucleotides (CPM). CPM were converted to bigwig files and visualized using Integrative Genomics Viewer (IGV) (Thorvaldsdóttir *et al*, 2013).

### QuantSeq data processing

#### Differential expression analysis

Differential expression for each pairwise comparison was assessed using the DESeq algorithm within the DESeq2 R package (Love *et al*, 2014). For each gene, this gives a base mean value, which is the mean of normalized counts for all samples after normalizing for sequencing depth, a value and standard error for the log2 fold change in relative RNA levels between the two samples being compared and a p-value, adjusted for multiple testing using the Benjamini-Hochberg method (padj), showing the significance of any difference.

#### Half-life comparison

Published genome-wide mRNA half-life data, determined using a metabolic labelling approach with the nucleoside analog 4-thiouridine (4SU) in HEK293T cells, were used (Lugowski *et al*, 2018). The log2 fold change in cytoplasmic RNA level for each gene was taken from the DESeq algorithm output for the comparison between cells kept at 37°C and cells transferred to 28°C, 18°C or 8°C for 24 h. In order to reduce noise and identify a relationship with mRNA half-life, data were processed as follows: Genes with missing half-life or log2 fold change values in any of the three temperature comparisons were excluded. Genes with extreme half-lives were excluded by removing those with a log2 half-life greater than two standard deviations away from the mean log2 half-life. For each cold temperature, genes were sorted by the log2 fold change for that temperature and split into 10 equal-sized groups. The mean log2 fold change and mean log2 half-life were then calculated for each group and plotted against each other. A linear regression line, Pearson correlation coefficient and p-value testing the significance of the correlation were calculated for each comparison.

#### Density scatter graphs comparing RNA level changes at different cold temperatures

Density scatter graphs were plotted showing the relationship in both the nucleus and cytoplasm between changes in RNA level upon transfer of cells from 37°C to 28°C or 18°C, 28°C or 8°C and 18°C or 8°C for 24h. This is done using the log2 fold change in RNA level for each gene from the DESeq algorithm output for the comparison between cells transferred to a specific cold temperature and cells kept at 37°C. Positive values show genes with higher relative RNA levels at the cold temperature. Genes with no padj or log2 fold change value in either of the two cold temperature v 37°C comparisons were excluded.

#### Density scatter graphs comparing nuclear and cytoplasmic RNA level changes

Density scatter graphs were plotted showing the relationship between changes in nuclear RNA levels and changes in cytoplasmic RNA levels from cells kept at 37°C and cells transferred from 37°C to a specific cold temperature for 5, 10 or 24 h. This is done using the log2 fold change in RNA level for each gene from the DESeq algorithm output for the comparison between nuclear fractions and for the comparison between cytoplasmic fractions. Positive values show genes with higher relative RNA levels at the cold temperature. Genes with a base mean value less than 30 in either the cytoplasmic or nuclear comparisons were excluded.

#### Standardized RNA level boxplots

RNA level counts for all genes for all samples from each time series were normalized by applying a regularized log transformation using the “rlogTransformation” function from the DESeq2 package with the blind option set to FALSE. The mean normalized count for each gene was then calculated across all repeats for each time point. The counts for each gene were then standardized so that the mean and standard deviation of the RNA level for each gene across the time-series was zero and one, respectively. This process makes the expression profiles of different transcripts across the time series comparable and was carried out separately for the nuclear and cytoplasmic fraction samples. Box and whisker plots displaying the range (whiskers), interquartile range (IQR, box edges), +/-IQR/[square root(number of genes)] (notch) and median (central line) of these standardized RNA levels were then plotted for nuclear and cytoplasmic fractions across each time series for subsets of genes selected according to whether they show significant (padj < 0.05) up or downregulation at a particular timepoint in the nuclear fraction.

#### GO term analysis

Genes showing significant (padj < 0.05) upregulation in the cytoplasm after exposing cells to 18°C for 24 h followed by 37°C for 2 h were submitted for gene-ontology (GO) analysis using the KEGG 2019 data base within Enrichr (Kuleshov *et al*, 2016).

#### Reads per million (RPM) bar graphs

Reads counts for each extended PAS were normalized for sequencing depth by dividing by the total number of PAS associated reads for each sample and then multiplying by 1 million to give reads per million (RPM). RPM were then obtained for each gene by combining the RPM for all PASs associated with each gene. RPM were then averaged across all replicates for each condition.

### Western Blot

1×10^6^ cells were trypsinized and washed with ice cold PBS. To assess whole cell protein levels, cell pellets were resuspended in 100 μl Lysis Buffer A, incubated (37°C, 20 min) with 0.5 μl Benzonase (Merck), and boiled (5 min) with 33 μl 4x Laemmli Buffer.

Protein samples were separated by gel electrophoresis on 10 % SDS polyacrylamide gels and transferred to nitrocellulose membranes using semi-dry transfer apparatus. Membranes were Ponceau stained, imaged and then washed in TBST (20 mM Tris-HCl (pH 7.5), 150 mM NaCl, 0.1 % Tween-20). Membranes were then incubated (4°C, overnight) with 5 % milk powder (BioRad) in TBST, then primary antibody in 5 % milk powder in TBST for 1.5 h, washed, then incubated with rabbit or mouse (Promega) HRP-conjugated secondary antibody diluted 1:4000 in 5 % milk powder in TBST for 45 min and washed again. Primary antibodies and their dilutions are detailed in Table S5. Antibody binding was visualized using chemiluminescence (Pierce) and X-ray film.

### RNA fluorescence in situ hybridisation (RNA-FISH)

#### RNA-FISH slide preparation

Cells were cultured on poly-L-lysine treated coverslips. Culture media was aspirated, and coverslips were washed once with PBS. Cells were fixed by incubating for 10 min with 4 % formaldehyde/PBS, washed twice with PBS, and permeabilized by incubating (>3 h, −20°C) in 70 % ethanol. Cells were rehydrated by incubating (5 min, RT) with FISH wash buffer (10 % formamide, 2x SSC). For hybridization, coverslips were placed cell-coated side down on a 48 μl drop containing 100 nM Quasar570-labelled probes complementary to one of *REV-ERBα, CRY2*, or *TP53* transcripts (Biosearch Technologies) (see Table S6 for probe sequences), 0.1 g/ml dextran sulfate, 1 mg/ml E. coli tRNA, 2 mM VRC, 20 μg/ml BSA, 2x SSC, 10 % formamide and incubated (37°C, 20 h) in a sealed parafilm chamber. Coverslips were twice incubated (37°C, 30 min) in pre-warmed FISH wash buffer, then in PBS containing 0.5 μg/ml 4’,6-diamidino-2-phenylindole (DAPI) (5 min, RT), washed twice with PBS, dipped in water, air-dried, placed cell-coated side down on a drop of ProLong Diamond Antifade Mountant (Life Technologies), allowed to polymerize for 24 h in the dark and then sealed with nail varnish.

#### RNA-FISH image acquisition and analysis

Cells were imaged using a DeltaVision Elite wide-field fluorescence deconvolution microscope using a 60x/1.40 objective lens (Olympus) and immersion oil with refractive index 1.514. First, the Quasar570-labelled RNA FISH probes were imaged using the TRITC channel, followed by imaging of the DAPI channel. The number of 0.2 μm-separated stacks imaged, number of replicates imaged for each condition and number of images across all combined replicates are listed in Table S7 (column “Stacks”, “Reps” and “Total images”, respectively). Images were deconvolved using the default conservative deconvolution method using DeltaVision Softworx software. Image quantification was carried out using Fiji (Schindelin *et al*, 2012). Deconvolved images were compressed to 2D images displaying the maximum intensity projection for each pixel across z-stacks listed in Table S7 (column “Projected”). Cell and nuclear areas were outlined using thresholding functions on the background TRITC signal and DAPI signal, respectively. Dots corresponding to transcripts were then counted for both nuclear and cytoplasmic areas for each image by applying the “Find Maxima” command with a noise tolerance specified in Table S7 (column “Maxima”). Bar charts show the mean number of dots per nuclear area and cytoplasmic area across all images for all combined replicates.

### Luciferase reporter assay

Two repeats of the six distinct profile experiment for both the 18°C for 5 h and 24 h incubations and 1 repeat of the four distinct profile experiment for an 18°C for 5 h incubation were carried out.

#### Six distinct profile experiment

In triplicate (i.e. 3 x 96 well plates), U2OS *PER2::LUC* normal (WT cells) and U2OS *PER2::LUC REV-ERBα* KO cells (KO cells) were each seeded at 20 % confluency in 30 wells of 96 well plates. After incubation for 24 h at 37°C, cells within a well were synchronized by incubating (45 min, 37°C) with 100 μl recording media (per litre: DMEM powder (Sigma cat#D2902), 1.9 g glucose, 10 mM HEPES, 1x GlutaMAX (ThermoFisher), 2.5 ml Penicillin/Steptomycin (Sigma cat#P0781), 4.7 ml NaHCO3 (7.5 % solution), final pH 7.3), containing 100 nM dexamethasone. Ten wells of each cell type in each plate were treated at one of three time intervals separated by 4 hours to give three groups of wells staggered at three different phases of the circadian cycle. After incubation, cells were washed twice with 100 μl prewarmed PBS and then 100 μl prewarmed recording media supplemented with 1x B-27 (ThermoFisher) and 100 μg/ml Luciferin was added. After all wells had been treated, all plates were sealed and incubated at 37°C in plate readers, whilst recording the bioluminescence at 30 min intervals. One control plate was kept at 37°C for at least 144 h and used as the control for both test plates. Test plates one and two were transferred to 18°C after 60 h and 72 h, respectively, after the start of the first dexamethasone treatment. This ensures that there are 6 groups of 10 wells of cells at 6 distinct phases of the circadian cycle at the point of 18°C exposure. Both test plates were returned to 37°C plate readers after either 5 h or 24 h and the bioluminescence was again recorded at 30 min intervals. Two repeats of this experiment were carried out for both 5 h and 24 h incubations. No control plate was recorded for the second repeat of the 5 h incubation.

#### Four distinct profile experiment

The same as for the six distinct profile experiment, except 15 wells of each cell type in each plate were treated at one of two time intervals separated by 12 hours to give two groups of wells staggered at two different phases of the circadian cycle. Test plates one and two were transferred to 18°C after 60 h and 66 h, respectively, after the start of the first dexamethasone treatment. This ensures that there are 4 groups of 15 wells of cells at 4 distinct phases of the circadian cycle at the point of 18°C exposure. Both test plates were returned to 37°C plate readers after 5 h. One repeat of this experiment was carried out.

### Luciferase reporter assay data processing

MultiCycle (Actimetrics) gives the period length and amplitude of recordings and these values were taken for all control wells across all control plates (plates kept at 37°C). The fold change in amplitude between WT and KO cells was calculated by dividing the amplitude of each well containing KO cells by that of the corresponding well in the same row containing WT cells within the same plate.

Two out of the total 540 individual well bioluminescence recordings across all experiments were discarded for technical errors during experimental set up. Raw bioluminescence recordings from all plates were baseline-detrended using MultiCycle. These were used to calculate the mean and standard deviation profiles for each group of synchronized plate wells. For all the six distinct profile experiments, all WT cell profiles were scaled to the same WT cell control mean profile so that their profile amplitudes for the period before the cold exposure were the same. All KO cell profiles were similarly scaled to the same KO cell control mean profile so that their profile amplitudes for the period before the cold exposure were also the same. The same scaling procedure was applied to the profiles from the four distinct profile experiment.

For scatter plots showing the change in phase variance resulting from 18°C incubations, for each well profile, the time to the second peak in bioluminescence after the start of the cold exposure and the equivalent peak in control well plates (this is the second peak after the end of the cold exposure for 24 h incubations) was determined. These times were then plotted for each condition and colored according to their distinct phase prior to 18°C exposure. This shows the effect of 18°C exposure on the spread of these profiles in comparison to profiles from cells kept at 37°C for both WT and KO cells. P-values (adjusted for multiple testing using the Holm-Bonferroni method) were calculated using the Brown-Forsythe method to test the significance of any difference in the variance for pairwise comparisons between 18°C-exposed cells and cells kept at 37°C and between WT and KO cells.

The same peak and the subsequent trough were used to calculate amplitude fold changes and phase shifts for profiles from cold-exposed wells relative to the corresponding mean control profile. Phase shifts as a percentage of period length were calculated using the following formula: ((peak time_cold-exposed_ + trough time_cold-exposed_)-(peak time_control_ + trough time_control_))/(2xmean period length). Mean shifts were calculated for all well profiles from each group with a distinct phase prior to 18°C exposure and for each experiment replicate. A one-sided, one sample t-test was then used to determine mean shifts significantly (p-value (adjusted for multiple testing using the Holm-Bonferroni method) < 0.05) greater than 5 % in either direction. Amplitude fold changes were calculated using the following formula (peak bioluminescence signal_cold-exposed_ – trough bioluminescence signal_cold-exposed_)/(peak bioluminescence signal_control_ – trough bioluminescence signal_control_). Mean log2 transformed fold changes were calculated for all well profiles from each group with a distinct phase prior to 18°C exposure and for each experiment replicate. A one-sided, one sample t-test was then used to determine mean changes significantly (p-value (adjusted for multiple testing using the Holm-Bonferroni method) < 0.05) greater or less than + or −, respectively, log2(1.4).

For experiments involving an 18°C for 5 h incubation, the position of the control profile at the point of rewarming within the circadian period of *PER2::LUC* expression was determined. As the phase of the REV-ERBα expression profile is shifted approximately 6 h in relation to PER2 expression in U2OS cells (Hoffmann *et al*, 2014), it is possible to predict the approximate level and direction of change of REV-ERBα expression at this point. Each mean phase shift and log2 amplitude fold change was plotted to approximately align with the position of its control profile at the point of rewarming within a circadian period represented by sine curves for *PER2:LUC* expression and predicted REV-ERBα expression.

### 3D-Structured Illumination Microscopy (3D-SIM)

#### 3D-SIM slide preparation

AC16 cells were cultured on 22×22 mm #1.5H high precision 170 μm ± 5 μm poly-L-lysine treated coverslips (Marienfeld Superior). After exposure to a particular temperature condition, culture media was aspirated, and coverslips were washed once with PBS. Cells were fixed by incubating for 10 min in 2% formaldehyde/PBS, then washed with 0.05 % Tween-20/PBS (PBST), permeabilized by incubating for 10 min in 0.2 % Triton-X-100/PBS, and then washed again with PBST. Coverslips were incubated (30 min, RT) in MaxBlock (ActiveMotif, cat#15252), then (overnight, 4°C) in MaxBlock containing primary antibodies against the proteins of interest (primary antibody details and dilutions are shown in Table S8), washed 4 times with PBST, then incubated (30 min, RT) in MaxBlock containing fluorescently labelled secondary antibodies (secondary antibody details and dilutions are shown in Table S8), washed again 4 times with PBST. Cells were post-fixed by incubating for 10 min in 4% formaldehyde/PBS and washed with PBST. Coverslips were incubated in PBST containing 2 μg/ml DAPI, washed with PBST, mounted in non-hardening Vectashield (Vector Laboratories, cat#H-1000) and stored at 4°C.

#### 3D-SIM image acquisition, reconstruction and quality control

3D-SIM image acquisition, reconstruction and quality control were carried out as detailed in Miron et al., 2019 (Miron *et al*, 2019). Images were acquired with a DeltaVision OMX V3 Blaze system (GE Healthcare) equipped with a 60x/1.42 NA PlanApo oil immersion objective (Olympus), pco.edge 5.5 sCMOS cameras (PCO) and 405, 488, 593 and 640 nm lasers. Spherical aberration was minimized using immersion oil with refractive index (RI) 1.514. 3D image stacks were acquired over the whole nuclear volume in z and with 15 raw images per plane (five phases, three angles). The raw data were computationally reconstructed with SoftWoRx 6.5.2 (GE Healthcare) using channel-specific OTFs recorded using immersion oil with RI 1.512, and Wiener filter settings between 0.002-0.006 to generate 3D stacks of 115 nm (488 nm) or 130 nm (593 nm) lateral and approximately 350 nm axial resolution. Multi-channel acquisitions were aligned in 3D using Chromagnon software (Matsuda *et al*, 2018) based on 3D-SIM acquisitions of multi-colour EdU-labelled C127 cells(Kraus *et al*, 2017). Table S9 details the number of nuclei imaged for each repeat, temperature condition and marker combination.

3D-SIM imaging and subsequent quantitative analyses are susceptible to artefacts (Demmerle *et al*, 2017). For instance, bulk labelling of densely packed chromatin inside mammalian nuclei of several μm depth entails high levels of out-of-focus blur, which reduces the contrast of illumination stripe modulation and thus the ability to recover high frequency (i.e., super-resolution) structure information. All SIM data were therefore routinely and meticulously quality controlled for effective resolution and absence of artifacts using SIMcheck (Ball *et al*, 2015), an open-source Image J plugin to assess SIM image quality via modulation contrast-to-noise ratio (MCNR), spherical aberration mismatch, reconstructed Fourier plot and reconstructed intensity histogram values (for more details see Demmerle et al., 2017 (Demmerle *et al*, 2017)).

To exclude potential false positive calls, we used the “MCNR map” macro of SIMcheck, which generates a metric of local stripe contrast in different regions of the raw data and directly correlates with the level of high-frequency information content in the reconstructed data (Demmerle *et al*, 2017). Only IF centroid signals with underlying MCNR values exceeding a stringent quality threshold were considered, while localisations with low underlying MCNR were discarded during the ChaiN analysis pipeline to exclude any SIM signal which falls below reconstruction confidence.

#### ChaiN - pipeline for high-content analysis of the 3D epigenome

ChaiN (for Chain high-throughput analysis of the in situ Nucleome), as detailed in Miron et al., 2019, consists of a pipeline of scripts for the automated high-throughput analyses presented in this investigation. Scripts are available from https://github.com/ezemiron/Chain. In brief, the DAPI-stain/chromatin channel is used to generate a nuclear mask and segment chromatin topology into 7 intensity/density classes using an R script that is expanding on a Hidden Markov model (Schmid *et al*, 2017). Class 1 denotes the interchromatin region while classes from 2-7 denote chromatin with increasing intensity/density. Based on this output of ChaiN, we quantitatively assessed the overall volume of chromatin (combined classes 2-7) versus the total nuclear volume (combined classes 1-7). Boxplots for nuclear volume and chromatin:nuclear volume display the median (central line), interquartile range (IQR, box edges) and most extreme data point no more than 1.5x IQR from box edges (whiskers). Outliers outside the whiskers are shown as individual points.

The IF signals for markers are thresholded by intensity using the Otsu algorithm, with all non-foci voxels replaced by zeros. The complement of this image is segmented by a 3D watershed algorithm and the 3D centroid coordinates extracted from each segmented region’s centre of mass after weighting by local pixel intensities. Potential false positive localizations were then filtered using the MCNR threshold detailed above to avoid potential artefacts skewing the statistical output. The quality controlled and filtered centroid positions were then related to the chromatin intensity/density classes. Centroid positions falling into each of the 7 segmented classes were counted, normalized to the respective class size and their relative enrichment (or depletion) displayed in a heat map on a log2-fold scale.

Nearest neighbor analysis was performed as follows. For each nucleus imaged, the Euclidian distance between each NPC centroid and all other hnRNPC centroids was calculated. The shortest of these distances was then taken and compiled into a list containing a distance for each NPC from all nuclei and from all temperature conditions. To remove outliers, distances greater than 2 standard deviations from the mean were excluded. Mean and S.E.M. distance was then calculated for each temperature condition.

## Supporting information

Movie chromatin classes at 8C supporting Figure 1A

Movie chromatin classes at 18C supporting Figure 1A

Movie chromatin classes at 28C supporting Figure 1A

Movie chromatin classes at 37C supporting Figure 1A

## Acknowledgments

We thank Paul Riley, Robin Choudhury, all members of the Furger and Mellor labs for helpful discussions and Nick Proudfoot, Shona Murphy and Andy Baldwin for comments on the manuscript. Funding: This research is funded by grants from the BBSRC awarded to AF (BB/N001184/1) and JM and AF (BB/S009035/1) and AJ (BB/N01992X/1 BBSRC, David Phillips fellowship). Imaging was performed at the Micron Oxford Advanced Bioimaging Unit funded by a Wellcome Trust Strategic Award (091911 and 107457/Z/15/Z).

## Author contributions

AF conceptualized and administered the project and supervised research planning. JM supervised RNA-FISH analysis. LS supervised 3D-SIM imaging experiments. AJ supervised the circadian rhythm experiments; HF was responsible for formal analysis, design and experimentation and wrote all in house-developed scripts (except 3D-SIM-related algorithms). RO and LS supported HF in 3D-SIM. DM supported HF in isolation and sequencing of RNA and analysis of RNA-Seq data. AF and HF wrote the manuscript with contributions from all other authors, HF created visualizations of the data with inputs from AF, JM, LS and AJ.

## Data availability

Underlying or raw data and/or further details of statistical analyses are available for Fig. 1B,D, 2B, 3B, 4E-H, 5D,E, S1D-I, S2A,C, S3F,G, S4A,B,F, S5B,C,E,F in the source data table. All RNA-Seq data are deposited at GEO (https://www.ncbi.nlm.nih.gov/geo/) under accession code GSE137003. All other data supporting the findings of this study are available from the corresponding authors on reasonable request.

## Supplementary Materials

<u>Supplementary Figures S1-S5</u>

<u>Supplementary Tables S1-S9</u>

<u>Fig. 5 extended data</u>

Table S1: Differential expression of core circadian clock and known cold or heat-induced gene transcripts in cells exposed to 18°C and subsequent rewarming.

Table S2: Gene ontology (GO) analysis of the cohort of transcripts that are upregulated in the cytoplasm of AC16 cells after rewarming.

Table S3: Oligonucleotides for *REV-ERBα* deletion and FLAG tagging

Table S4: Number of RNA-Seq sample replicates for each condition

Table S5: Western blot primary antibodies and their dilutions.

Table S6: RNA-FISH probes

Table S7: RNA-FISH image acquisition

Table S8: Antibodies used in 3D-SIM analysis

Table S9: 3D-SIM image acquisition

Movies: Temperature induces changes to the higher-order chromatin architecture imaged through the nuclear volume

## References

Al-Fageeh MB & Smales CM (2006) Control and regulation of the cellular responses to cold shock: The responses in yeast and mammalian systems. Biochem. J. 397: 247–259 Available at: http://www.biochemj.org/content/397/2/247.abstract

Ann Ran F, Hsu PD, Wright J, Agarwala V, Scott DA & Zhang F (2013) Genome engineering using the CRIPR-Cas9 system. Nature

Ball G, Demmerle J, Kaufmann R, Davis I, Dobbie IM & Schermelleh L (2015) SIMcheck: a Toolbox for Successful Super-resolution Structured Illumination Microscopy. Sci. Rep. 5: 15915 Available at: https://doi.org/10.1038/srep15915

Bastide A, Peretti D, Knight JRP, Grosso S, Spriggs R V., Pichon X, Sbarrato T, Roobol A, Roobol J, Vito D, Bushell M, von der Haar T, Smales CM, Mallucci GR & Willis AE (2017) RTN3 Is a Novel Cold-Induced Protein and Mediates Neuroprotective Effects of RBM3. Curr. Biol. 27: 638–650 Available at: http://linkinghub.elsevier.com/retrieve/pii/S0960982217300829 [Accessed March 17, 2017]

Bautista DM, Siemens J, Glazer JM, Tsuruda PR, Basbaum AI, Stucky CL, Jordt S-E & Julius D (2007) The menthol receptor TRPM8 is the principal detector of environmental cold. Nature 448: 204–208 Available at: https://doi.org/10.1038/nature05910

Bootman MD, Fearnley C, Smyrnias I, Macdonald F & Roderick HL (2009) An update on nuclear calcium signalling.

Brajkovic D & Ducharme MB (2006) Facial cold-induced vasodilation and skin temperature during exposure to cold wind. Eur. J. Appl. Physiol. 96: 711–721

Brown SA, Zumbrunn G, Fleury-Olela F, Preitner N & Schibler U (2002) Rhythms of Mammalian Body Temperature Can Sustain Peripheral Circadian Clocks. Curr. Biol. 12: 1574–1583 Available at: https://www.sciencedirect.com/science/article/pii/S0960982202011454 [Accessed July 11, 2018]

Buhr ED, Yoo SH & Takahashi JS (2010) Temperature as a universal resetting cue for mammalian circadian oscillators. Science (80-.). 330: 379–385 Available at: http://science.sciencemag.org/content/330/6002/379.abstract

Calixto CPG, Guo W, James AB, Tzioutziou NA, Entizne JC, Panter PE, Knight H, Nimmo HG, Zhang R & Brown JWS (2018) Rapid and dynamic alternative splicing impacts the arabidopsis cold response transcriptome[CC-BY]. Plant Cell 30: 1424–1444

Cho H, Zhao X, Hatori M, Yu RT, Barish GD, Lam MT, Chong L-W, DiTacchio L, Atkins AR, Glass CK, Liddle C, Auwerx J, Downes M, Panda S & Evans RM (2012) Regulation of circadian behaviour and metabolism by REV-ERB-α and REV-ERB-β. Nature 485: 123 Available at: https://doi.org/10.1038/nature11048

Choukèr A, Bereiter-Hahn J, Singer D & Heldmaier G (2019) Hibernating astronauts---science or fiction? Pflügers Arch. - Eur. J. Physiol. 471: 819–828 Available at: https://doi.org/10.1007/s00424-018-2244-7

Chua BA, Van Der Werf I, Jamieson C & Signer RAJ (2020) Post-Transcriptional Regulation of Homeostatic, Stressed, and Malignant Stem Cells. Cell Stem Cell 26: 138–159 Available at: https://doi.org/10.1016/j.stem.2020.01.005

Davidson MM, Nesti C, Palenzuela L, Walker WF, Hernandez E, Protas L, Hirano M & Isaac ND (2005) Novel cell lines derived from adult human ventricular cardiomyocytes. J. Mol. Cell. Cardiol. 39: 133–147

Demmerle J, Innocent C, North AJ, Ball G, Müller M, Miron E, Matsuda A, Dobbie IM, Markaki Y & Schermelleh L (2017) Strategic and practical guidelines for successful structured illumination microscopy. Nat. Protoc. 12: 988 Available at: https://doi.org/10.1038/nprot.2017.019

Ding Y, Shi Y & Yang S (2019) Advances and challenges in uncovering cold tolerance regulatory mechanisms in plants. New Phytol. 222: 1690–1704

Giuliodori AM, Di Pietro F, Marzi S, Masquida B, Wagner R, Romby P, Gualerzi CO & Pon CL (2010) The cspA mRNA Is a Thermosensor that Modulates Translation of the Cold-Shock Protein CspA. Mol. Cell 37: 21–33 Available at: http://dx.doi.org/10.1016/j.molcel.2009.11.033

Gordon CJ (2001) The therapeutic potential of regulated hypothermia. Emerg. Med. J. 18: 81 LP – 89 Available at: http://emj.bmj.com/content/18/2/81.abstract

Hattori K, Ishikawa H, Sakauchi C, Takayanagi S, Naguro I & Ichijo H (2017) Cold stress-induced ferroptosis involves the ASK1-p38 pathway. EMBO Rep. 18: 2067 LP – 2078 Available at: http://embor.embopress.org/content/18/11/2067.abstract

Hoffmann J, Symul L, Shostak A, Fischer T, Naef F & Brunner M (2014) Non-Circadian Expression Masking Clock-Driven Weak Transcription Rhythms in U2OS Cells. PLoS One 9: e102238

Jackson TC & Kochanek PM (2019) A New Vision for Therapeutic Hypothermia in the Era of Targeted Temperature Management: A Speculative Synthesis. Ther. Hypothermia Temp. Manag. 9: 13–47 Available at: https://doi.org/10.1089/ther.2019.0001

Jagannath A, Butler R, Godinho SIH, Couch Y, Brown LA, Vasudevan SR, Flanagan KC, Anthony D, Churchill GC, Wood MJA, Steiner G, Ebeling M, Hossbach M, Wettstein JG, Duffield GE, Gatti S, Hankins MW, Foster RG & Peirson SN (2013) The CRTC1-SIK1 Pathway Regulates Entrainment of the Circadian Clock. Cell 154: 1100–1111 Available at: https://www.sciencedirect.com/science/article/pii/S0092867413009616?via%3Dihub [Accessed October 25, 2019]

Jagannath A, Taylor L, Wakaf Z, Vasudevan SR & Foster RG (2017) The genetics of circadian rhythms, sleep and health. Hum. Mol. Genet. 26: R128–R138 Available at: https://doi.org/10.1093/hmg/ddx240

Kandror O, Bretschneider N, Kreydin E, Cavalieri D & Goldberg AL (2004) Yeast Adapt to Near-Freezing Temperatures by STRE/Msn2,4-Dependent Induction of Trehalose Synthesis and Certain Molecular Chaperones. Mol. Cell 13: 771–781 Available at: https://www.sciencedirect.com/science/article/pii/S1097276504001480?via%3Dihub [Accessed July 17, 2018]

Kim J-M, Sasaki T, Ueda M, Sako K & Seki M (2015) Chromatin changes in response to drought, salinity, heat, and cold stresses in plants. Front. Plant Sci. 6: 114 Available at: http://journal.frontiersin.org/article/10.3389/fpls.2015.00114

Kim YH, Marhon SA, Zhang Y, Steger DJ, Won K-J & Lazar MA (2018) Rev-erbα dynamically modulates chromatin looping to control circadian gene transcription. Science (80-.). 359: 1274 LP – 1277 Available at: http://science.sciencemag.org/content/359/6381/1274.abstract

Koike N, Yoo S-H, Huang H-C, Kumar V, Lee C, Kim T-K & Takahashi JS (2012) Transcriptional Architecture and Chromatin Landscape of the Core Circadian Clock in Mammals. Science (80-.). 338: 349 LP – 354 Available at: http://science.sciencemag.org/content/338/6105/349.abstract

Kon N, Yoshikawa T, Honma S, Yamagata Y, Yoshitane H, Shimizu K, Sugiyama Y, Hara C, Kameshita I & Honma K (2014) CaMKII is essential for the cellular clock and coupling between morning and evening behavioral rhythms. : 1101–1110

Kornmann B, Schaad O, Bujard H, Takahashi JS & Schibler U (2007) System-Driven and Oscillator-Dependent Circadian Transcription in Mice with a Conditionally Active Liver Clock. PLOS Biol. 5: e34 Available at: https://doi.org/10.1371/journal.pbio.0050034

Kraus F, Miron E, Demmerle J, Chitiashvili T, Budco A, Alle Q, Matsuda A, Leonhardt H, Schermelleh L & Markaki Y (2017) Quantitative 3D structured illumination microscopy of nuclear structures. Nat. Protoc. 12: 1011 Available at: https://doi.org/10.1038/nprot.2017.020

Kuleshov M V, Jones MR, Rouillard AD, Fernandez NF, Duan Q, Wang Z, Koplev S, Jenkins SL, Jagodnik KM, Lachmann A, McDermott MG, Monteiro CD, Gundersen GW & Ma’ayan A (2016) Enrichr: a comprehensive gene set enrichment analysis web server 2016 update. Nucleic Acids Res. 44: W90–W97 Available at: http://dx.doi.org/10.1093/nar/gkw377

Kültz D (2020) Evolution of cellular stress response mechanisms. J. Exp. Zool. Part A Ecol. Integr. Physiol.: 1–20

Kutcher ME, Forsythe RM & Tisherman SA (2016) Emergency preservation and resuscitation for cardiac arrest from trauma. Int. J. Surg. 33: 209–212 Available at: http://dx.doi.org/10.1016/j.ijsu.2015.10.014

Liu X, Freitas J, Zheng D, Oliveira MS, Hoque M, Martins T, Henriques T, Tian B & Moreira A (2017) Transcription elongation rate has a tissue-specific impact on alternative cleavage and polyadenylation in Drosophila melanogaster. RNA 23: 1807–1816

Liu Y, Hu W, Murakawa Y, Yin J, Wang G, Landthaler M & Yan J (2013) Cold-induced RNA-binding proteins regulate circadian gene expression by controlling alternative polyadenylation. Sci Rep 3: 2054 Available at: http://www.ncbi.nlm.nih.gov/pubmed/23792593 [Accessed September 5, 2013]

Londo JP, Kovaleski AP & Lillis JA (2018) Divergence in the transcriptional landscape between low temperature and freeze shock in cultivated grapevine (Vitis vinifera). Hortic. Res. 5: 10 Available at: https://doi.org/10.1038/s41438-018-0020-7

Love MI, Huber W & Anders S (2014) Moderated estimation of fold change and dispersion for RNA-seq data with DESeq2. Genome Biol. 15: 550

Lugowski A, Nicholson B & Rissland OS (2018) DRUID: a pipeline for transcriptome-wide measurements of mRNA stability. RNA 24: 623–632

Mackensen GB, McDonagh DL & Warner DS (2009) Perioperative Hypothermia: Use and Therapeutic Implications. J. Neurotrauma 26: 342–358 Available at: https://doi.org/10.1089/neu.2008.0596

Matsuda A, Schermelleh L, Hirano Y, Haraguchi T & Hiraoka Y (2018) Accurate and fiducial-marker-free correction for three-dimensional chromatic shift in biological fluorescence microscopy. Sci. Rep. 8: 7583 Available at: https://doi.org/10.1038/s41598-018-25922-7

Miron E, Oldenkamp R, Pinto DMS, Brown JM, Faria ARC, Shaban HA, Rhodes JDP, Innocent C, de Ornellas S, Buckle V & Schermelleh L (2019) Chromatin arranges in filaments of blobs with nanoscale functional zonation. bioRxiv: 566638 Available at: http://biorxiv.org/content/early/2019/03/04/566638.abstract

Montaigne D, Marechal X, Modine T, Coisne A, Mouton S, Fayad G, Ninni S, Klein C, Ortmans S, Seunes C, Potelle C, Berthier A, Gheeraert C, Piveteau C, Deprez R, Eeckhoute J, Duez H, Lacroix D, Deprez B, Jegou B, et al (2018) Daytime variation of perioperative myocardial injury in cardiac surgery and its prevention by Rev-Erbα antagonism: a single-centre propensity-matched cohort study and a randomised study. Lancet 391: 59–69 Available at: https://www.sciencedirect.com/science/article/pii/S0140673617321323?via%3Dihub [Accessed June 13, 2018]

Morf J, Rey G, Schneider K, Stratmann M, Fujita J, Naef F & Schibler U (2012) Cold-Inducible RNA-Binding Protein Modulates Circadian Gene Expression Posttranscriptionally. Science (80-.). 338: 379 LP – 383 Available at: http://science.sciencemag.org/content/338/6105/379.abstract

Nagano AJ, Kawagoe T, Sugisaka J, Honjo MN, Iwayama K & Kudoh H (2019) Annual transcriptome dynamics in natural environments reveals plant seasonal adaptation. Nat. Plants 5: 74–83 Available at: https://doi.org/10.1038/s41477-018-0338-z

Neve J, Burger K, Li W, Hoque M, Patel R, Tian B, Gullerova M & Furger A (2016) Subcellular RNA profiling links splicing and nuclear DICER1 to alternative cleavage and polyadenylation. Genome Res. 26: 24–35 Available at: http://genome.cshlp.org/lookup/doi/10.1101/gr.193995.115

Nicetto D & Zaret KS (2019) Role of H3K9me3 heterochromatin in cell identity establishment and maintenance. Curr. Opin. Genet. Dev. 55: 1–10 Available at: https://www.sciencedirect.com/science/article/pii/S0959437X19300127?via%3Dihub [Accessed May 23, 2019]

Nordeen CA & Martin SL (2019) Engineering Human Stasis for Long-Duration Spaceflight. Physiology 34: 101–111 Available at: https://doi.org/10.1152/physiol.00046.2018

Ou J, Ball JM, Luan Y, Zhao T, Miyagishima KJ, Xu Y, Zhou H, Chen J, Merriman DK, Xie Z, Mallon BS & Li W (2018) iPSCs from a Hibernator Provide a Platform for Studying Cold Adaptation and Its Potential Medical Applications. Cell 173: 851–863.e16 Available at: https://www.sciencedirect.com/science/article/pii/S0092867418302903?via%3Dihub [Accessed July 18, 2018]

Pariollaud M, Gibbs JE, Hopwood TW, Brown S, Begley N, Vonslow R, Poolman T, Guo B, Saer B, Jones DH, Tellam JP, Bresciani S, Tomkinson NCO, Wojno-Picon J, Cooper AWJ, Daniels DA, Trump RP, Grant D, Zuercher W, Willson TM, et al (2018) Circadian clock component REV-ERBα controls homeostatic regulation of pulmonary inflammation. J. Clin. Invest. 128: 2281–2296 Available at: https://doi.org/10.1172/JCI93910

Park J, Lim CJ, Shen M, Park HJ, Cha J-Y, Iniesto E, Rubio V, Mengiste T, Zhu J-K, Bressan RA, Lee SY, Lee B-H, Jin JB, Pardo JM, Kim W-Y & Yun D-J (2018) Epigenetic switch from repressive to permissive chromatin in response to cold stress. Proc. Natl. Acad. Sci. 115: E5400 LP–E5409 Available at: http://www.ncbi.nlm.nih.gov/pubmed/29784800 [Accessed July 20, 2018]

Peretti D, Bastide A, Radford H, Verity N, Molloy C, Martin MG, Moreno JA, Steinert JR, Smith T, Dinsdale D, Willis AE & Mallucci GR (2015) RBM3 mediates structural plasticity and protective effects of cooling in neurodegeneration. Nature 518: 236–239 Available at: http://dx.doi.org/10.1038/nature14142

Phengchat R, Takata H, Morii K, Inada N, Murakoshi H, Uchiyama S & Fukui K (2016) Calcium ions function as a booster of chromosome condensation. Sci. Rep. 6: 1–10

Pires R, Jones NS, Andreu L, Gupta R, Enver T & Francisco J (2010) Connecting Variability in Global Transcription Rate to Mitochondrial Variability. 8:

Quinones QJ, Ma Q, Zhang Z, Barnes BM & Podgoreanu M V (2014) Organ Protective Mechanisms Common to Extremes of Physiology: A Window through Hibernation Biology. Integr. Comp. Biol. 54: 497–515 Available at: http://dx.doi.org/10.1093/icb/icu047

Rao V, Feindel CM, Cohen G, Borger MA, Boylen P & Ross HJ (2001) Is profound hypothermia required for storage of cardiac allografts? J. Thorac. Cardiovasc. Surg. 122: 501–507 Available at: https://www.sciencedirect.com/science/article/pii/S0022522301771747?via%3Dihub [Accessed July 18, 2018]

Richardson CD, Ray GJ, DeWitt MA, Curie GL & Corn JE (2016) Enhancing homology-directed genome editing by catalytically active and inactive CRISPR-Cas9 using asymmetric donor DNA. Nat. Biotechnol.

Ruf T & Geiser F (2015) Daily torpor and hibernation in birds and mammals. Biol. Rev. 90: 891–926

Rzechorzek NM, Connick P, Patani R, Selvaraj BT & Chandran S (2015) Hypothermic Preconditioning of Human Cortical Neurons Requires Proteostatic Priming. EBioMedicine 2: 528–535 Available at: http://linkinghub.elsevier.com/retrieve/pii/S2352396415000985 [Accessed August 29, 2017]

Schermelleh L, Carlton PM, Haase S, Shao L, Winoto L, Kner P, Burke B, Cardoso MC, Agard DA, Gustafsson MGL, Leonhardt H & Sedat JW (2008) Subdiffraction Multicolor Imaging of the Nuclear Periphery with 3D Structured Illumination Microscopy. Science (80-.). 320: 1332 LP – 1336 Available at: http://science.sciencemag.org/content/320/5881/1332.abstract

Schindelin J, Arganda-Carreras I, Frise E, Kaynig V, Longair M, Pietzsch T, Preibisch S, Rueden C, Saalfeld S, Schmid B, Tinevez J-Y, White DJ, Hartenstein V, Eliceiri K, Tomancak P & Cardona A (2012) Fiji: an open-source platform for biological-image analysis. Nat. Methods 9: 676–682

Schmid VJ, Cremer M & Cremer T (2017) Quantitative analyses of the 3D nuclear landscape recorded with super-resolved fluorescence microscopy. Methods 123: 33–46 Available at: https://www.sciencedirect.com/science/article/pii/S1046202316303322?via%3Dihub [Accessed October 25, 2019]

Sulli G, Rommel A, Wang X, Kolar MJ, Puca F, Saghatelian A, Plikus M V, Verma IM & Panda S (2018) Pharmacological activation of REV-ERBs is lethal in cancer and oncogene-induced senescence. Nature 553: 351 Available at: http://dx.doi.org/10.1038/nature25170

Takahashi JS (2017) Transcriptional architecture of the mammalian circadian clock. Nat. Rev. Genet. 18: 164–179 Available at: http://dx.doi.org/10.1038/nrg.2016.150

Tamaru T, Hattori M, Honda K, Benjamin I, Ozawa T & Takamatsu K (2011) Synchronization of Circadian Per2 Rhythms and HSF1-BMAL1:CLOCK Interaction in Mouse Fibroblasts after Short-Term Heat Shock Pulse. PLoS One 6: e24521 Available at: https://doi.org/10.1371/journal.pone.0024521

Thorvaldsdóttir H, Robinson JT & Mesirov JP (2013) Integrative Genomics Viewer (IGV): High-performance genomics data visualization and exploration. Brief. Bioinform. 14: 178–192

Vargas DY, Raj A, Marras SAE, Kramer FR & Tyagi S (2005) Mechanism of mRNA transport in the nucleus. 102:

Vihervaara A, Duarte FM & Lis JT (2018) Molecular mechanisms driving transcriptional stress responses. Nat. Rev. Genet. 19: 385–397 Available at: http://dx.doi.org/10.1038/s41576-018-0001-6

Wang R, Nambiar R, Zheng D & Tian B (2018) PolyA_DB 3 catalogs cleavage and polyadenylation sites identified by deep sequencing in multiple genomes. Nucleic Acids Res. 46: D315–D319

Yin Y, Lu JY, Zhang X, Shao W, Xu Y, Li P, Hong Y, Cui L, Shan G, Tian B, Zhang QC & Shen X (2020) U1 snRNP regulates chromatin retention of noncoding RNAs. Nature 580: 147–150 Available at: http://dx.doi.org/10.1038/s41586-020-2105-3

Zeng Z, Zhang W, Marand AP, Zhu B, Buell CR & Jiang J (2019) Cold stress induces enhanced chromatin accessibility and bivalent histone modifications H3K4me3 and H3K27me3 of active genes in potato. Genome Biol. 20: 1–17

Zhang Y, Burkhardt DH, Rouskin S, Li G-W, Weissman JS & Gross CA (2018) A Stress Response that Monitors and Regulates mRNA Structure Is Central to Cold Shock Adaptation. Mol. Cell 70: 274–286.e7 Available at: https://www.sciencedirect.com/science/article/pii/S1097276518301801 [Accessed June 19, 2018]

Zhang Y, Fang B, Emmett MJ, Damle M, Sun Z, Feng D, Armour SM, Remsberg JR, Jager J, Soccio RE, Steger DJ & Lazar MA (2015) GENE REGULATION. Discrete functions of nuclear receptor Rev-erbα couple metabolism to the clock. Science 348: 1488–1492 Available at: https://www.ncbi.nlm.nih.gov/pubmed/26044300

Zhao C, Lang Z & Zhu J-K (2015) Cold responsive gene transcription becomes more complex. Trends Plant Sci. 20: 466–468 Available at: https://www.sciencedirect.com/science/article/pii/S1360138515001478 [Accessed July 17, 2018]

Zhao X, Hirota T, Han X, Cho H, Chong L-W, Lamia K, Liu S, Atkins AR, Banayo E, Liddle C, Yu RT, Yates JR, Kay SA, Downes M & Evans RM (2016) Circadian Amplitude Regulation via FBXW7-Targeted REV-ERBα Degradation. Cell 165: 1644–1657 Available at: https://www.sciencedirect.com/science/article/pii/S0092867416305608?via%3Dihub#! [Accessed March 26, 2019]

Zhu J-K (2016) Abiotic Stress Signaling and Responses in Plants. Cell 167: 313–324 Available at: https://www.sciencedirect.com/science/article/pii/S0092867416310807?via%3Dihub [Accessed July 18, 2018]

Zhu X, Bührer C & Wellmann S (2016) Cold-inducible proteins CIRP and RBM3, a unique couple with activities far beyond the cold. Cell. Mol. Life Sci. 73: 3839–59 Available at: http://dx.doi.org/10.1007/s00018-016-2253-7

Zhuang X, Magri A, Hill M, Lai AG, Kumar A, Rambhatla SB, Donald CL, Lopez-Clavijo AF, Rudge S, Pinnick K, Chang WH, Wing PAC, Brown R, Qin X, Simmonds P, Baumert TF, Ray D, Loudon A, Balfe P, Wakelam M, et al (2019) The circadian clock components BMAL1 and REV-ERBα regulate flavivirus replication. Nat. Commun. 10: 377 Available at: https://doi.org/10.1038/s41467-019-08299-7

