## Supplementary tables and figures for "Cold induced chromatin compaction and nuclear retention of clock mRNAs resets the circadian rhythm"

**Affiliations:**

**This PDF fsection includes:**

Figs. S1 to S5, Fig. 5 extended data
Tables S1 to S9

**Table S1.**

Differential expression of core circadian clock and known cold or heat-induced gene transcripts in cells exposed to 18°C and subsequent rewarming.

| Gene | 37°C v 18°C 24h | Adjusted p-value | 37°C v 18°C 24h, then 37°C 2h | Adjusted p-value | 37°C v 18°C 24h, then 37°C 5h | Adjusted p-value |
|---|---|---|---|---|---|---|
| REV-ERBα | 4.001** | 2.19E-18 | 2.461** | 4.09E-06 | 0.135 | NA |
| PER1 | 0.263 | 0.921 | 1.622** | 0.021 | 0.058 | 0.970 |
| PER2 | 1.071 | 0.516 | 3.323** | 7.22E-12 | 2.285 | NA |
| CRY1 | 0.381 | 0.810 | 0.994 | 0.158 | 1.883* | 0.084 |
| CRY2 | -0.067 | 0.974 | 1.559** | 4.93E-04 | 0.074 | 0.950 |
| ARNTL | 0.052 | 0.991 | 0.266 | 0.829 | -0.075 | 0.972 |
| CLOCK | -0.335 | 0.899 | 0.133 | 0.911 | 0.270 | 0.864 |
| TP53 | -0.364 | 0.791 | -0.330 | 0.630 | -0.608 | 0.529 |
| RBM3 | 0.394 | 0.777 | 0.118 | 0.885 | -0.240 | 0.811 |
| CIRBP | 0.430 | 0.672 | 0.171 | 0.817 | -0.378 | 0.667 |
| DNAJA1 | 0.182 | 0.933 | 0.096 | 0.915 | 0.377 | 0.736 |
| DNAJA2 | 0.170 | 0.927 | 0.349 | 0.532 | 0.466 | 0.585 |
| DNAJA3 | -0.790 | 0.383 | -0.993 | 0.112 | -0.664 | 0.452 |
| DNAJA4 | 0.520 | 0.785 | 0.412 | 0.685 | 0.310 | NA |
| DNAJB1 | -0.482 | 0.743 | -0.600 | 0.331 | 0.004 | 0.998 |
| DNAJB11 | 0.195 | 0.926 | 0.262 | 0.754 | 0.275 | 0.811 |
| DNAJB12 | -0.503 | 0.633 | -0.237 | 0.698 | -0.152 | 0.875 |
| DNAJB13 | -0.387 | NA | 1.564 | NA | -0.224 | NA |
| DNAJB14 | 0.630 | 0.866 | -0.393 | NA | 0.501 | NA |
| DNAJB2 | -0.467 | 0.622 | -0.110 | 0.844 | -0.476 | 0.481 |
| DNAJB4 | -0.017 | 0.992 | 0.104 | 0.921 | 0.962 | 0.412 |
| DNAJB5 | 0.040 | 0.990 | -0.651 | 0.407 | -1.270 | 0.332 |
| DNAJB6 | 0.103 | 0.962 | 0.102 | 0.890 | 0.257 | 0.789 |
| DNAJB9 | -0.515 | 0.752 | 0.432 | 0.622 | 0.277 | 0.821 |
| DNAJC1 | -0.063 | 0.987 | 0.129 | 0.914 | 0.890 | 0.520 |
| DNAJC10 | 0.254 | 0.894 | 0.447 | 0.557 | 0.973 | 0.359 |
| DNAJC11 | -0.110 | 0.959 | 0.072 | 0.926 | -0.178 | 0.847 |
| DNAJC12 | -0.392 | 0.922 | 0.890 | NA | 0.772 | NA |
| DNAJC13 | 0.058 | 0.991 | 0.711 | 0.420 | 0.178 | 0.913 |
| DNAJC14 | 0.028 | 0.992 | 0.241 | 0.836 | -1.168 | 0.452 |
| DNAJC15 | -0.534 | 0.821 | 0.301 | 0.727 | 0.227 | 0.865 |
| DNAJC16 | 0.490 | 0.804 | -0.147 | 0.907 | 0.355 | NA |

| | | | | | | |
|---|---|---|---|---|---|---|
| DNAJC17 | -1.000 | 0.697 | -0.153 | 0.925 | 0.098 | NA |
| DNAJC18 | 0.160 | 0.936 | -0.264 | 0.730 | -0.252 | 0.828 |
| DNAJC19 | -0.171 | 0.938 | -0.056 | 0.954 | -0.414 | 0.666 |
| DNAJC2 | 0.444 | 0.817 | 0.367 | 0.716 | 0.954 | 0.603 |
| DNAJC21 | 0.265 | 0.881 | 0.722 | 0.282 | 0.344 | 0.756 |
| DNAJC22 | 0.299 | 0.930 | 0.440 | 0.685 | 0.254 | NA |
| DNAJC24 | -0.113 | 0.981 | -0.858 | NA | 0.059 | NA |
| DNAJC25 | 0.245 | 0.931 | -0.236 | 0.857 | 0.534 | NA |
| DNAJC27 | -1.090 | 0.729 | -0.097 | NA | -0.194 | NA |
| DNAJC28 | 0.857 | 0.846 | -0.475 | NA | -1.158 | NA |
| DNAJC3 | 0.556 | 0.802 | 1.271 | 0.158 | 1.780 | 0.221 |
| DNAJC30 | -0.340 | 0.742 | -0.374 | 0.520 | -1.121 | 0.203 |
| DNAJC4 | -0.661 | 0.756 | -0.359 | 0.781 | -1.124 | NA |
| DNAJC5 | -0.531 | 0.540 | -0.383 | 0.399 | -0.524 | 0.478 |
| DNAJC5B | 1.468 | NA | 0.943 | NA | 1.357 | NA |
| DNAJC6 | -0.149 | 0.971 | 0.356 | 0.762 | 0.298 | NA |
| DNAJC7 | -0.025 | 0.992 | -0.756 | 0.648 | -0.326 | 0.850 |
| DNAJC8 | -0.012 | 0.993 | -0.248 | 0.691 | -0.094 | 0.924 |
| DNAJC9 | 0.469 | 0.684 | 0.852 | 0.142 | 0.686 | 0.452 |
| HIKESHI | -0.085 | 0.978 | 0.196 | 0.843 | 0.496 | 0.701 |
| HSBP1 | 0.017 | 0.991 | 0.008 | 0.988 | -0.041 | 0.963 |
| HSBP1L1 | 0.500 | 0.872 | 0.291 | NA | 1.325 | NA |
| HSF1 | 0.163 | 0.952 | -0.690 | 0.213 | -0.548 | 0.513 |
| HSF2 | -0.019 | 0.992 | 1.017 | 0.128 | 0.199 | 0.869 |
| HSF2BP | -0.420 | 0.876 | -0.140 | 0.917 | -0.288 | NA |
| HSF4 | 0.548 | 0.866 | -0.354 | NA | -1.106 | NA |
| HSF5 | 0.819 | NA | 0.369 | NA | -0.727 | NA |
| HSP90AA1 | 0.168 | 0.948 | 0.697 | 0.371 | 0.890 | 0.438 |
| HSP90AB1 | 0.289 | 0.824 | 0.211 | 0.768 | 0.216 | 0.830 |
| HSP90B1 | 0.432 | 0.808 | 0.265 | 0.763 | 0.452 | 0.703 |
| HSPA12A | 0.600 | 0.758 | 0.452 | 0.568 | -0.393 | 0.772 |
| HSPA12B | 0.105 | NA | 0.952 | NA | 2.123 | NA |
| HSPA13 | 0.551 | 0.681 | 0.290 | 0.703 | 0.781 | 0.438 |
| HSPA14 | 0.052 | 0.980 | 0.651 | 0.195 | 0.666 | 0.388 |
| HSPA1A | -0.260 | 0.883 | -0.427 | 0.534 | 0.140 | 0.895 |
| HSPA1B | -0.477 | 0.746 | -0.071 | 0.944 | 0.381 | 0.749 |
| HSPA1L | 1.456 | NA | 1.446 | NA | 1.719 | NA |
| HSPA2 | -0.836 | 0.463 | -0.021 | 0.982 | -0.928 | 0.402 |

| | | | | | | |
|---|---|---|---|---|---|---|
| HSPA4 | 0.042 | 0.987 | 0.495 | 0.408 | 0.758 | 0.398 |
| HSPA4L | 0.136 | 0.979 | -0.476 | NA | 0.338 | NA |
| HSPA5 | 0.659 | 0.671 | 0.561 | 0.489 | 0.635 | 0.598 |
| HSPA8 | -0.537 | 0.740 | 0.068 | 0.945 | -0.156 | 0.902 |
| HSPA9 | 0.437 | 0.721 | 0.506 | 0.376 | 0.684 | 0.449 |
| HSPB1 | -0.109 | 0.964 | 0.512 | 0.591 | -0.348 | 0.748 |
| HSPB11 | 0.223 | 0.872 | 0.366 | 0.562 | 0.503 | 0.534 |
| HSPB2 | -0.261 | 0.954 | -1.816 | NA | -3.315 | NA |
| HSPB3 | 0.404 | 0.837 | 0.996 | 0.150 | -0.277 | 0.847 |
| HSPB6 | -0.303 | 0.892 | -0.646 | 0.390 | -1.103 | 0.411 |
| HSPB7 | 1.730 | NA | 0.760 | NA | 1.589 | NA |
| HSPB8 | 0.660 | 0.811 | 1.521 | 0.129 | 1.645 | NA |
| HSPB9 | -0.387 | NA | 1.127 | NA | -0.224 | NA |
| HSPD1 | 0.053 | 0.982 | 0.211 | 0.769 | 0.859 | 0.378 |
| HSPE1 | -0.080 | 0.963 | 0.610 | 0.323 | 0.453 | 0.563 |
| HSPH1 | 0.316 | 0.850 | 0.279 | 0.714 | 1.023 | 0.297 |

Differential expression analysis carried out using the DESeq algorithm within the DESeq2 R package (Love et al., 2014) for AC16 cells exposed to the temperature conditions shown for core circadian clock genes (*REV-ERBα*, *PER1*, *PER2*, *CRY1*, *CRY2*, *CLOCK*, *ARNTL*), known cold-induced genes (*CIRBP*, *RBM3*) and known heat shock-induced genes (all other listed genes). The log2 fold change in expression for each gene for each temperature condition comparison is shown with positive values indicating genes upregulated in the latter condition. P-values, adjusted for multiple testing using the Benjamini-Hochberg method show the significance of each change. ** = change with adjusted p-value < 0.05, * = change with adjusted p-value < 0.1. DESeq output sets some p-values to NA when the mean normalized count is low.

**Table S2.**

Gene ontology (GO) analysis of the cohort of transcripts that are upregulated in the cytoplasm of AC16 cells after rewarming.

| Term | Overlap | P-value | Adjusted p-value | Genes |
|---|---|---|---|---|
| Circadian rhythm | 5/31 | 8.31E-07 | 0.000256 | PER2; PER1; BHLHE40; CRY2; REV-ERBα |
| Transcriptional misregulation in cancer | 5/186 | 0.00427 | 0.658 | PER2; SIX4; SIN3A; JMJD1C; KLF3 |
| p53 signaling pathway | 3/72 | 0.00808 | 0.830 | APAF1; SIAH1; PMAIP1 |
| Apoptosis | 4/143 | 0.00911 | 0.701 | BCL2L11; APAF1; PMAIP1; CFLAR |
| Parathyroid hormone synthesis secretion and action | 3/106 | 0.0227 | 1 | GNA13; AKAP13; HBEGF |

Gene ontology (GO) analysis of the cohort of transcripts that are upregulated in the cytoplasm of AC16 cells transferred to 18°C for 24h and then rewarmed to 37°C for 2h compared to cells kept at 37°C. The top five terms with lowest p-values. Only the term “circadian rhythm” is significantly enriched (adjusted p-value < 0.05) after adjustment for multiple testing using the Benjamini-Hochberg method. GO analysis was performed using the Enrichr (Kuleshov et al., 2016) platform against the KEGG 2019 Human data base of GO pathways.

**Table S3**

Oligonucleotides for *REV-ERBα* deletion and FLAG tagging

| sgRNA | Oligonucleotide 1 | Oligonucleotide 2 |
|---|---|---|
| Tag | CACCGtggacgcccagtgacccgcc | AAACggcgggtcactgggcgtccaC |
| Del US | CACCGtagtccaccgacaaagtggg | AAACccccactttgtcggtggactaC |
| Del DS | CACCGttagcaaatctccgggccga | AAACtcggcccggagatttgctaaC |

Tagging ssODN:
CTGGTTTGCTTTTCCTTTTCGTCTCGTAAAGGAGAGAGAAGTGCAGAGTTCGATTCTG
TACAAGGGGGCAGCGGCAGAAGGCCGGCCGGGCGGGTCACTTGTCATCGTCATCCT
TGTAATCGATATCATGATCTTTATAATCACCGTCATGGTCTTTGTAGTCCTGGGCGTC
CACCCGGAAGGACAGCAGCTTCTCGGAA

**Table S4**

RNA-Seq samples for each condition and replicates

| Cell Line | Condition | Spiked | Cyt or Nuc | Seq type | No. of Repeats |
|---|---|---|---|---|---|
| AC16 | 37°C | No | Cytoplasm | QuantSeq | 6 |
| AC16 | 37°C | No | Nucleus | QuantSeq | 7 |
| AC16 | 28°C 24h | No | Cytoplasm | QuantSeq | 6 |
| AC16 | 28°C 24h | No | Nucleus | QuantSeq | 6 |
| AC16 | 28°C 24h, 37°C 2h | No | Cytoplasm | QuantSeq | 2 |
| AC16 | 28°C 24h, 37°C 2h | No | Nucleus | QuantSeq | 2 |
| AC16 | 18°C 5h | No | Cytoplasm | QuantSeq | 4 |
| AC16 | 18°C 5h | No | Nucleus | QuantSeq | 4 |
| AC16 | 18°C 10h | No | Cytoplasm | QuantSeq | 4 |
| AC16 | 18°C 10h | No | Nucleus | QuantSeq | 4 |
| AC16 | 18°C 24h | No | Cytoplasm | QuantSeq | 6 |
| AC16 | 18°C 24h | No | Nucleus | QuantSeq | 6 |
| AC16 | 18°C 24h, 37°C 2h | No | Cytoplasm | QuantSeq | 4 |
| AC16 | 18°C 24h, 37°C 2h | No | Nucleus | QuantSeq | 4 |
| AC16 | 18°C 24h, 37°C 5h | No | Cytoplasm | QuantSeq | 2 |
| AC16 | 18°C 24h, 37°C 5h | No | Nucleus | QuantSeq | 2 |
| AC16 | 18°C 24h, 37°C 10h | No | Cytoplasm | QuantSeq | 2 |
| AC16 | 18°C 24h, 37°C 10h | No | Nucleus | QuantSeq | 2 |
| AC16 | 18°C 24h, 37°C 24h | No | Cytoplasm | QuantSeq | 2 |
| AC16 | 18°C 24h, 37°C 24h | No | Nucleus | QuantSeq | 2 |
| AC16 | 8°C 24h | No | Cytoplasm | QuantSeq | 6 |
| AC16 | 8°C 24h | No | Nucleus | QuantSeq | 6 |
| AC16 | 37°C | Yes | Cytoplasm | QuantSeq | 4 |
| AC16 | 37°C | Yes | Nucleus | QuantSeq | 4 |
| AC16 | 18°C 24h | Yes | Cytoplasm | QuantSeq | 4 |
| AC16 | 18°C 24h | Yes | Nucleus | QuantSeq | 4 |
| AC16 | 18°C 24h, 37°C 2h | Yes | Cytoplasm | QuantSeq | 4 |
| AC16 | 18°C 24h, 37°C 2h | Yes | Nucleus | QuantSeq | 4 |
| U2OS | 37°C | No | Cytoplasm | QuantSeq | 2 |
| U2OS | 37°C | No | Nucleus | QuantSeq | 2 |
| U2OS | 18°C 5h | No | Cytoplasm | QuantSeq | 2 |
| U2OS | 18°C 5h | No | Nucleus | QuantSeq | 2 |
| U2OS | 18°C 24h | No | Cytoplasm | QuantSeq | 2 |
| U2OS | 18°C 24h | No | Nucleus | QuantSeq | 2 |
| U2OS | 18°C 24h, 37°C 2h | No | Cytoplasm | QuantSeq | 2 |
| U2OS | 18°C 24h, 37°C 2h | No | Nucleus | QuantSeq | 2 |
| AC16 | 37°C | No | Cytoplasm | Full-length | 1 |
| AC16 | 37°C | No | Nucleus | Full-length | 1 |
| AC16 | 18°C 24h | No | Cytoplasm | Full-length | 1 |
| AC16 | 18°C 24h | No | Nucleus | Full-length | 1 |

All samples sequenced and the number of biological replicates.

**Table S5**

Western blot primary antibodies and their dilutions.

| **Primary antibody (Dilution)** | **Source** |
|---|---|
| Mouse mAb anti-Flag (1:4000) | Sigma F3165, clone M2 |
| Rabbit pAb anti-TP53 (1:2000) | Bethyl A300-247A |

**Table S6**

RNA-FISH probes

| Probe | Sequences |
|---|---|
| *REV-ERBα* | gtgcaaaagtcccagagga, aaggtagcaaggagggtcg, agagagagtgtgtaggggg, tctgctttgcatgggaaga, gttggacgttgaggcaacg, acgatcaggatccgaagca, aactagaggttgcgatcgc, ggtcattcaaactggacct, atgtcttcaccagctgaga, tgttgttgttggagtccag, actggagccaatgtaggtg, atagagggattcagggctg, ttgggtcagggactggaag, ggatggtgggaagtaggtg, tggaatgctcccaaaggag, catggccacttgtagactc, gatgttgctggtgctcttg, gtaacaccatgccattcag, aacgtccccacacacttta, gtgcacaccgtagtggaag, ctggatgttctgctggatg, ttgcgattgatgcggacga, ttgaagcgacattgctggc, agacatgcccacagagaga, cttctctcgtttggggatg, cactctgcatctcagcaag, gctgttgtggaaactggga, tcatgggcgtaggtgaaga, ttgaagttgccaggtgagc, ctacctgatgcatggttgg, catttagggcctcgttatg, cgttgctgttggactggtg, aacattctttgagttgccc, catgcgggtacatgttcat, tttggcaaactctaccacc, gtgacttggtcatgctgag, caaaggtgccagccttaag, aagcaaagcgcaccatcag, ggtccttcacgttgaacaa, ggcttaggaacatcactgt, gaacatggcactgagcagg, gttgagcttctcgctgaag, ctgcagagacaagcaccac, gaagcggaattctccatgc, cggttcttcagcaccagag, agcttggtgaagcgggaag, atgcatgttgttcagggtc, gaaggacagcagcttctcg, |
| *CRY2* | cttccagagactgaagtagg, gtttccttaaacttgtgtcc, actacaaacaggcgggagtt, tgaacagccttgggaacacg, catattcaaaggtcaagcgg, ttctttcccaaagggttcag, agaattctccgtcactactt, aatgatcctgtccaggtcat, cctgaaagcgcttgtatgta, ttccgttccaagtgcttatc, ggtctctcatagttggcaac, gcaggagagacaaccaaagc, cacaggcggtagtagaagag, gcttcacctttttatacagg, caaatagggagaggggaggt, aagaactctcgccataggag, aaacctggggttgttggtag, aggggatctggatgcagatg, tggcatcaatccaagggaag, ttcaatttgggcaggtatcg, agggctcatagatgtatcga, cttctgaattgactctgggg, cacaccaatgatgcacttgg, tctcggcatggttgacgatg, ttcgttcaatgttaagccgg, cgcgaaagctgctggtaaat, aaacagcactggcgtgctac, aaagtcactgccataacctc, cttgcaggaacaggtctcag, cactacgttctgttcagaca, aatcccatgctttggacaac, tacgtctggacagcatccat, ttagatctgtctatggcctc, tctgtagggtggtggaaact, cagtcaggtatctgtgtgta, cttaggctcctattacagta, agcttctggggaaggaacaa, agtgagtcagtttgaccctt, ccctggaagccaacagaata, agtctggatgctgacaagtc, ctactgggatagctgacatg, gttctgggtgtaatctctac, tactgctcctggaaggaatg, attggcttctctgggtcaaa |
| *TP53* | gtgtcaccgtcgtggaaag, catggcagtgacccggaag, tcgacgctaggatctgact, gggggacagaacgttgttt, ttcaatatcgtccggggac, gtaggttttctgggaaggg, cttggccagttggcaaaac, taagatgctgaggaggggc, tgtagttgtagtggatggt, ttctttggctggggagagg, ccacggatctgaagggtga, gtagactgacccttttgg, atgtcagtctgagtcaggc, agcaagggttcaaagaccc, cttctgacgcacacctatt, caacttgttcagtggagcc, atctaagctggtatgtcct, tccctcacagtaaaaacct, catttcttacatctcccaa, actaacccttaactgcaag, cctacctagaatgtggctg, cctggttagtacggtgaag, tcaacagtgagggacagct, gttctagaccccatgtaat, aacaagcaccctcaagggg, caccgaccaacagggagag, gctgcccaactgtagaaac, acaactccctctacctaac, aggttgtcagacagggttt, taggtactaaggttcacca, tgggatggggtgagatttc, gagatgaaatcctccaggg, ggtggatccagatcatcat, ccctgagcataaaacaagt, atgcagatgtgcttgcaga |

**Table S7**
RNA-FISH image acquisition

| Probe | Cell Line | Condition | Reps | Total Images | Stacks | Projected | Maxima |
|---|---|---|---|---|---|---|---|
| *REV-ERBα* | AC16 | 37°C | 4 | 65 | 30 | 5-25 | 700 |
| *REV-ERBα* | AC16 | 18°C 5h | 2 | 30 | 30 | 5-25 | 700 |
| *REV-ERBα* | AC16 | 18°C 5h, 37°C 2h | 2 | 21 | 30 | 5-25 | 700 |
| *REV-ERBα* | AC16 | 18°C 24h | 4 | 63 | 30 | 5-25 | 700 |
| *REV-ERBα* | AC16 | 18°C 24h, 37°C 2h | 4 | 64 | 30 | 5-25 | 700 |
| *REV-ERBα* | U2OS | 37°C | 3 | 61 | 40 | 8-33 | 700 |
| *REV-ERBα* | U2OS | 18°C 5h | 3 | 61 | 40 | 8-33 | 700 |
| *REV-ERBα* | U2OS | 18°C 5h, 37°C 2h | 3 | 69 | 40 | 8-33 | 700 |
| *REV-ERBα* | U2OS | 18°C 24h | 3 | 53 | 40 | 8-33 | 700 |
| *REV-ERBα* | U2OS | 18°C 24h, 37°C 2h | 3 | 65 | 40 | 8-33 | 700 |
| *CRY2* | AC16 | 37°C | 2 | 43 | 20 | 3-17 | 6000 |
| *CRY2* | AC16 | 18°C 24h | 2 | 42 | 20 | 3-17 | 6000 |
| *CRY2* | AC16 | 18°C 24h, 37°C 2h | 2 | 42 | 20 | 3-17 | 6000 |
| *TP53* | AC16 | 37°C | 3 | 47 | 30 | 4-26 | 1600 |
| *TP53* | AC16 | 8°C 24h | 3 | 45 | 30 | 4-26 | 1600 |
| *TP53* | AC16 | 18°C 24h, 37°C 2h | 3 | 48 | 30 | 4-26 | 1600 |

Total images = Number of images across all replicates
Stacks = Number of 0.2 μm stacks imaged
Projected = Stacks used for maximum intensity projection
Maxima = Noise tolerance set for the “Find Maxima” algorithm

**Table S8**

Antibodies used in 3D-SIM analysis

| **Primary antibody (Dilution)** | **Source** |
|---|---|
| Rat mAb anti-Pol2S2P (1:1000) | Millipore 041571, Clone 3E10 |
| Rabbit pAb anti-H3K4me3 (1:1000) | ActiveMotif 39159 |
| Mouse mAb anti-H3K27me3 (1:500) | Abcam ab6002, clone mAbcam6002 |
| Mouse mAb anti-H3K9me3 (1:500) | ActiveMotif 61013, clone MABI0319 |
| Rabbit mAb anti-hnRNPC1/C2 (1:500) | Abcam ab133607, clone EPNCIR152 |
| Mouse mAb anti-nuclear pore complex (NPC) (1:1000) | Abcam ab24700, QE5 |
| **Secondary antibody (All 1:500 dilution)** | **Source** |
| Alexa-488 Goat pAb anti-Rabbit-IgG | ThermoFisher A11029 |
| Alexa-594 Donkey pAb anti-Mouse-IgG | ThermoFisher A21203 |
| Alexa-488 Goat pAb anti-Rat-IgG | ThermoFisher A11006 |

**Table S9**
3D-SIM image acquisition

| Condition | Alexa-488 Marker | Alexa-594 Marker | Repeat | No. of Cells Imaged |
|---|---|---|---|---|
| 37°C | HNRNPC | NPC | 1 | 31 |
| 28°C 24h | HNRNPC | NPC | 1 | 26 |
| 18°C 5h | HNRNPC | NPC | 1 | 19 |
| 18°C 24h | HNRNPC | NPC | 1 | 22 |
| 18°C 24h, 37°C 2h | HNRNPC | NPC | 1 | 25 |
| 8°C 24h | HNRNPC | NPC | 1 | 19 |
| 37°C | HNRNPC | NPC | 2 | 12 |
| 37°C | H3K4me3 | H3K27me3 | 2 | 13 |
| 37°C | Pol2S2P | H3K9me3 | 2 | 17 |
| 28°C 24h | HNRNPC | NPC | 2 | 11 |
| 28°C 24h | H3K4me3 | H3K27me3 | 2 | 12 |
| 28°C 24h | Pol2S2P | H3K9me3 | 2 | 13 |
| 18°C 5h | HNRNPC | NPC | 2 | 10 |
| 18°C 5h | H3K4me3 | H3K27me3 | 2 | 12 |
| 18°C 5h | Pol2S2P | H3K9me3 | 2 | 20 |
| 18°C 24h | HNRNPC | NPC | 2 | 12 |
| 18°C 24h | H3K4me3 | H3K27me3 | 2 | 12 |
| 18°C 24h | Pol2S2P | H3K9me3 | 2 | 12 |
| 18°C 24h, 37°C 2h | HNRNPC | NPC | 2 | 12 |
| 18°C 24h, 37°C 2h | H3K4me3 | H3K27me3 | 2 | 12 |
| 18°C 24h, 37°C 2h | Pol2S2P | H3K9me3 | 2 | 11 |
| 8°C 24h | HNRNPC | NPC | 2 | 12 |
| 8°C 24h | H3K4me3 | H3K27me3 | 2 | 12 |
| 8°C 24h | Pol2S2P | H3K9me3 | 2 | 12 |

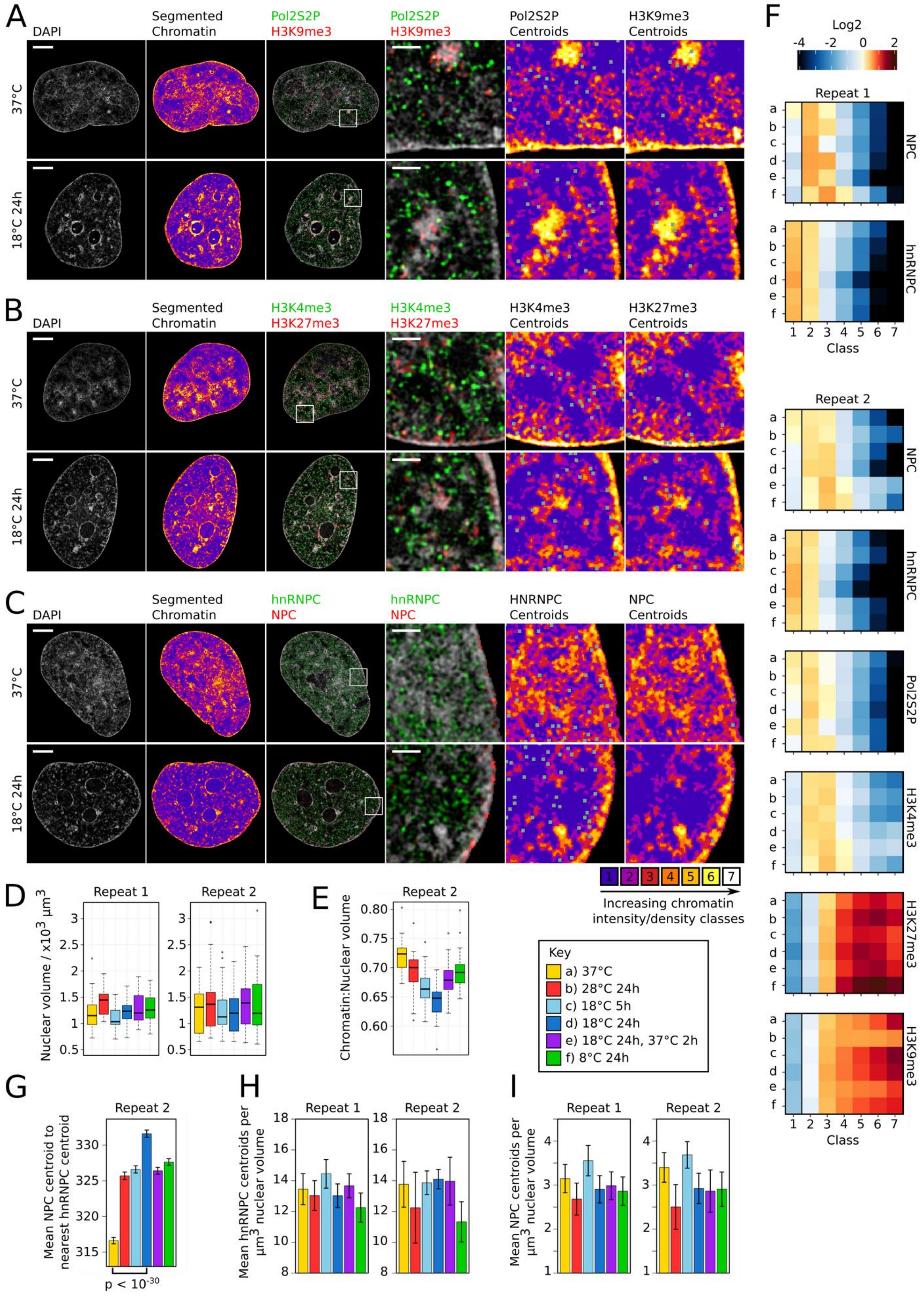

A
DAPI
Segmented Chromatin
Pol2S2P H3K9me3
Pol2S2P H3K9me3
Pol2S2P Centroids
H3K9me3 Centroids
37°C
18°C 24h
B
DAPI
Segmented Chromatin
H3K4me3 H3K27me3
H3K4me3 H3K27me3
H3K4me3 Centroids
H3K27me3 Centroids
37°C
18°C 24h
C
DAPI
Segmented Chromatin
hnRNPC NPC
hnRNPC NPC
HNRNPC Centroids
NPC Centroids
37°C
18°C 24h
1 2 3 4 5 6 7
Increasing chromatin intensity/density classes
D
Repeat 1
Repeat 2
Nuclear volume / x10$^3$ µm$^3$
E
Repeat 2
Chromatin:Nuclear volume
Key
a) 37°C
b) 28°C 24h
c) 18°C 5h
d) 18°C 24h
e) 18°C 24h, 37°C 2h
f) 8°C 24h
F
Log2
Repeat 1
NPC
hnRNPC
Class
Repeat 2
NPC
hnRNPC
Pol2S2P
H3K4me3
H3K27me3
H3K9me3
Class
G
Repeat 2
Mean NPC centroid to nearest hnRNPC centroid
p < 10$^{-30}$
H
Repeat 1
Repeat 2
Mean hnRNPC centroids per µm$^3$ nuclear volume
I
Repeat 1
Repeat 2
Mean NPC centroids per µm$^3$ nuclear volume

**Fig. S1**

**A-C**) Representative single z-planes of 3D-SIM image stacks of the DAPI-stained nuclei (column 1, 3 and 4) of AC16 cells kept at 37°C or exposed to 18°C for 24h showing the spatial distribution of the IF signal of nuclear markers (column 3 and 4): serine 2-phosphorylated RNA polymerase II (Pol2S2P, green) and H3K9me3 (red) (A); H3K4me3 (green) and H3K27me3 (red) (B); hnRNPC (green) and nuclear pore complexes (NPCs, red) (C). Segmented chromatin images (column 2, 5 and 6) show the DAPI signal segmented into 7 different chromatin classes according to relative intensity. The lowest (class 1) denotes the interchromatin (IC) region while classes from 2 to 7 denote regions with increasing chromatin densities. Classes 2-7 have been grouped together as one region in Fig. 1A. These are overlaid with green pixels with a grey outline indicating marker centroid coordinates for centroids located in the z-plane shown (column 5 and 6). Scale bar whole nuclei: 10 μm. Scale bar enlarged section: 1 μm. **D**) Boxplots of the nuclear volume for cells exposed to different temperature conditions for biological repeats 1 and 2. **E**) Boxplots of the ratio of chromatin volume to nuclear volume for cells exposed to different temperature conditions from a second biological repeat. DAPI signal is segmented into chromatin and IC regions according to relative intensity. **F**) Heatmaps of the log2 fold change in IF signal relative to a random distribution for each of the 7 chromatin density classes for all temperature conditions (a-f (see Key)) and for each marker for biological repeats 1 and 2. **G**) Bar graph of the mean NPC centroid to nearest hnRNPC centroid distance for each temperature condition for a second biological repeat. Error bars show the S.E.M. p-value (two-sided Welch's t-test) calculated for the comparison shown. **H** and **I**) Bar graph of the mean hnRNPC (H) or NPC (I) centroids per $\mu m^3$ nuclear volume for all cells imaged for biological repeats 1 and 2. Error bars show the standard deviation.

### Figure S2

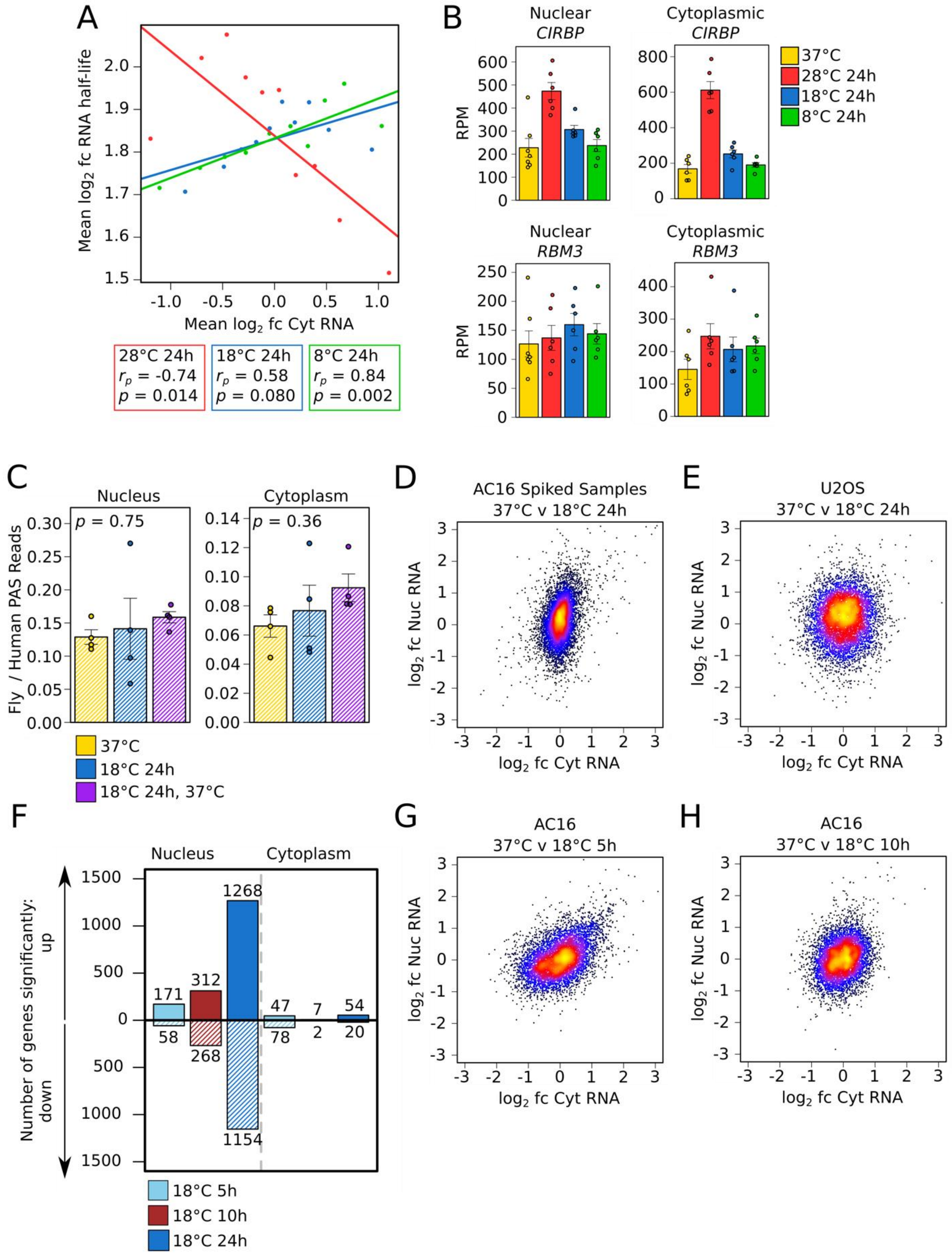
A
Mean log2 fc RNA half-life
Mean log2 fc Cyt RNA
28°C 24h
rp = -0.74
p = 0.014
18°C 24h
rp = 0.58
p = 0.080
8°C 24h
rp = 0.84
p = 0.002
B
Nuclear CIRBP
Cytoplasmic CIRBP
Nuclear RBM3
Cytoplasmic RBM3
RPM
37°C
28°C 24h
18°C 24h
8°C 24h
C
Nucleus
Cytoplasm
p = 0.75
p = 0.36
Fly / Human PAS Reads
37°C
18°C 24h
18°C 24h, 37°C
D
AC16 Spiked Samples
37°C v 18°C 24h
log2 fc Nuc RNA
log2 fc Cyt RNA
E
U2OS
37°C v 18°C 24h
F
Nucleus
Cytoplasm
Number of genes significantly:
up
down
171
312
1268
47
7
54
58
268
1154
78
2
20
18°C 5h
18°C 10h
18°C 24h
G
AC16
37°C v 18°C 5h
H
AC16
37°C v 18°C 10h

**Fig S2.**

**A**) Relationship between mean RNA half-life taken from (Lugowski et al., 2018) and mean log2 fold change in cytoplasmic RNA level in cells exposed to 28°C (red), 18°C (blue), or 8°C (green) for 24h compared to cells kept at 37°C for genes that have been ordered by fold change in cytoplasmic RNA level and then binned into 10 equal-sized groups. Linear regression line, Pearson correlation coefficient ($r_p$), and p-value (p) testing the significance of the correlation are shown for each comparison. **B**) Mean nuclear and cytoplasmic RNA levels in reads per million (RPM) of known cold-induced genes *CIRBP* and *RBM3* for cells kept at 37°C or transferred from 37°C to 28°C, 18°C, or 8°C for 24h. Error bars show S.E.M. Dots show the value for each biological repeat. **C**) Mean ratio of *D. melanogaster* (fly) aligned polyA site (PAS) reads to human aligned PAS reads for nuclear and cytoplasmic samples prepared from AC16 cells harvested after exposure to the three different conditions shown and spiked with a known ratio of *D. melanogaster* Schneider 2 (S2) cells. p-values (one-way ANOVA) testing the significance of the difference between means are shown. Error bars show S.E.M. Dots show the value for each biological repeat. n = 4. As these ratios do not show significant changes, a change in the RNA level of a gene upon exposure to 18°C and/or upon subsequent rewarming represents a change in its absolute RNA level. **D**-**E**) Relationship between the log2 fold change in RNA level in the nucleus and that in the cytoplasm for all genes upon transfer of cells from 37°C to 18°C for 24h in AC16 cells (*D. melanogaster* spiked in samples) (D) or U2OS cells (E). **F**) Number of genes showing significant (adjusted $p < 0.05$) up or downregulation in nuclear or cytoplasmic RNA levels upon transfer of cells from 37°C to 18°C for 5, 10 or 24h. **G**-**H**) Relationship between the log2 fold change in RNA level in the nucleus and that in the cytoplasm for all genes upon transfer of cells from 37°C to 18°C for 5h (G) or 10h (H). Points (D-E and G-H) are color-coded from low to high density (black < blue < red < yellow).

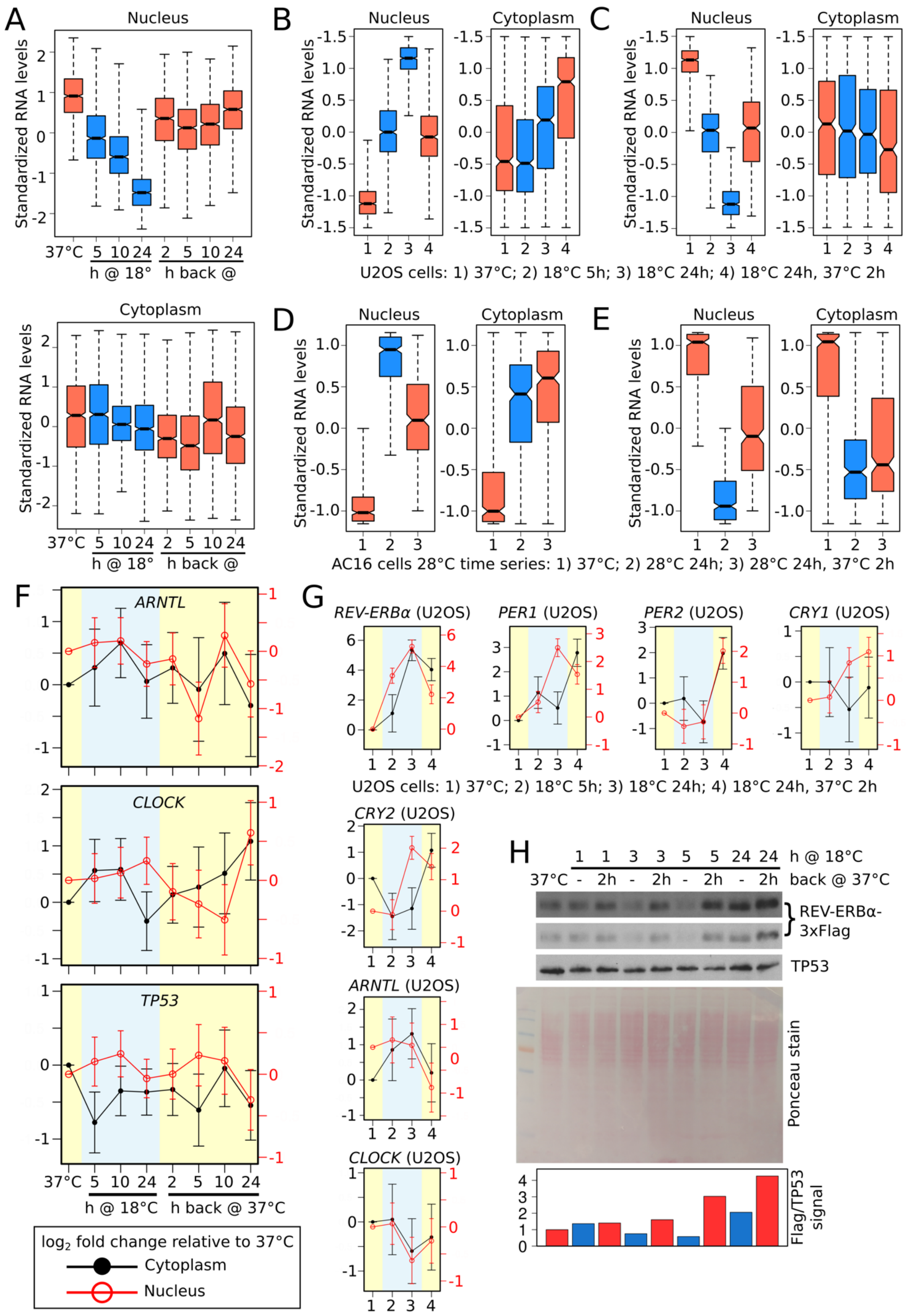
A
Nucleus
Standardized RNA levels
37°C
5 10 24
h @ 18°
2 5 10 24
h back @
Cytoplasm
B
Nucleus
Cytoplasm
C
Nucleus
Cytoplasm
U2OS cells: 1) 37°C; 2) 18°C 5h; 3) 18°C 24h; 4) 18°C 24h, 37°C 2h
D
Nucleus
Cytoplasm
E
Nucleus
Cytoplasm
AC16 cells 28°C time series: 1) 37°C; 2) 28°C 24h; 3) 28°C 24h, 37°C 2h
F
ARNTL
CLOCK
TP53
37°C 5 10 24 2 5 10 24
h @ 18°C
h back @ 37°C
log2 fold change relative to 37°C
Cytoplasm
Nucleus
G
REV-ERBα (U2OS)
PER1 (U2OS)
PER2 (U2OS)
CRY1 (U2OS)
U2OS cells: 1) 37°C; 2) 18°C 5h; 3) 18°C 24h; 4) 18°C 24h, 37°C 2h
CRY2 (U2OS)
ARNTL (U2OS)
CLOCK (U2OS)
H
1 1 3 3 5 5 24 24 h @ 18°C
37°C - 2h - 2h - 2h - 2h back @ 37°C
REV-ERBα-3xFlag
TP53
Ponceau stain
Flag/TP53 signal

**Fig. S3**

**A**) Boxplots of standardized nuclear and cytoplasmic RNA levels at time points during the transfer of AC16 cells from 37°C to 18°C for 24h and then back to 37°C for 24h for the group of 1154 genes showing significant downregulation in nuclear RNA levels at the 18°C for 24h time point. **B** and **C**) Boxplots of standardized nuclear and cytoplasmic RNA levels at time points during the transfer of U2OS cells from 37°C to 18°C for 24h and then back to 37°C for 2h for the group of 815 genes showing significant upregulation (B) or 552 genes showing significant downregulation (C) in nuclear RNA levels at the 18°C for 24h time point. **D**) and **E**) Boxplots of standardized nuclear and cytoplasmic RNA levels at time points during the transfer of AC16 cells from 37°C to 28°C for 24h and then back to 37°C for 2h for the group of 304 genes showing significant upregulation (D) or 347 genes showing significant downregulation (E) in nuclear RNA levels at the 28°C for 24h time point. **F**) Log2 fold change in cytoplasmic (black line, left axis) and nuclear (red line, right axis) RNA level of core circadian clock activator genes *ARNTL* (*BMAL1*) and *CLOCK*, and the control gene *TP53* at time points during the transfer of AC16 cells from 37°C to 18°C for 24h and then back to 37°C for 24h relative to cells kept at 37°C. **G**) Log2 fold change in cytoplasmic (black line, left axis) and nuclear (red line, right axis) RNA level of core circadian clock genes at time points during the transfer of U2OS cells from 37°C to 18°C for 24h and then back to 37°C for 2h relative to cells kept at 37°C. Error bars (F and G) show the standard error. **H**) Western blot of Flag-tagged REV-ERBα levels (short and long exposure) in AC16 cells transferred from 37°C to 18°C for the time periods indicated and also returned after each of these time periods to 37°C for 2h (biological repeat of Western blot in Fig. 3D). TP53 was used as a loading control as its transcript levels show minimal changes in response to 18°C exposure (Fig. 4, S3F). Ponceau stain is shown as an additional loading control. Quantification of the REV-ERBα/TP53 signal is shown below.

### Figure S4

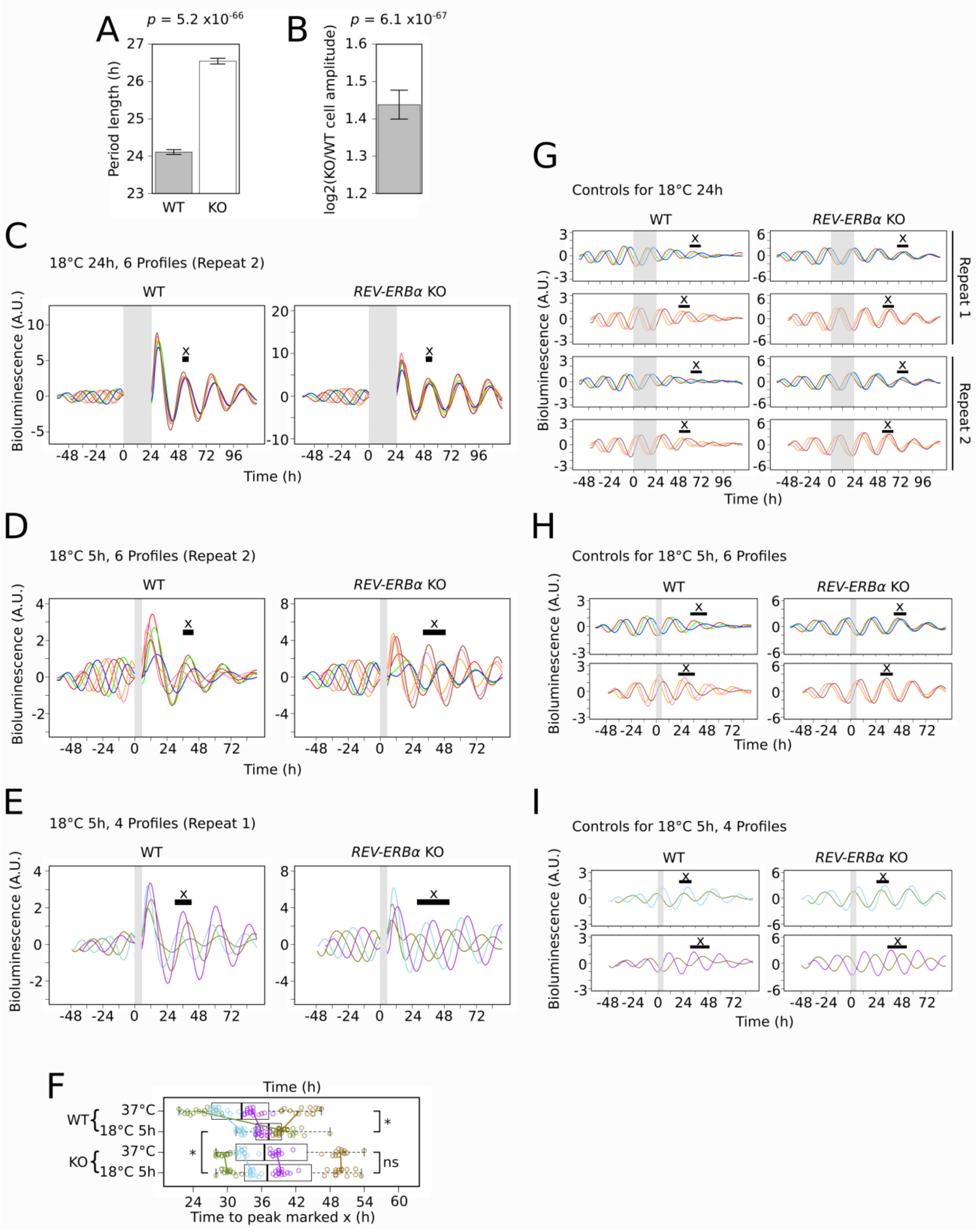

A
p = 5.2 x10-66
Period length (h)
WT
KO
B
p = 6.1 x10-67
log2(KO/WT cell amplitude)
C
18°C 24h, 6 Profiles (Repeat 2)
WT
REV-ERBα KO
Bioluminescence (A.U.)
Time (h)
D
18°C 5h, 6 Profiles (Repeat 2)
WT
REV-ERBα KO
Bioluminescence (A.U.)
Time (h)
E
18°C 5h, 4 Profiles (Repeat 1)
WT
REV-ERBα KO
Bioluminescence (A.U.)
F
Time (h)
WT
37°C
18°C 5h
KO
37°C
18°C 5h
ns
Time to peak marked x (h)
G
Controls for 18°C 24h
WT
REV-ERBα KO
Repeat 1
Repeat 2
Bioluminescence (A.U.)
Time (h)
H
Controls for 18°C 5h, 6 Profiles
WT
REV-ERBα KO
Bioluminescence (A.U.)
Time (h)
I
Controls for 18°C 5h, 4 Profiles
WT
REV-ERBα KO
Bioluminescence (A.U.)
Time (h)

**Fig. S4**

**A**) Bar graph showing the mean period length of WT and *REV-ERBα* KO cells. P-value from a two-sided Welch's t-test to determine the significance of the difference of the means. **B**) Bar graph showing the mean log2(*REV-ERBα* KO/WT cell amplitude). Error bars show the S.E.M. P-value from a two-sided, one sample t-test to determine the significance of the difference of the mean from 0. Values were calculated from the period length and amplitude derived using MultiCycle for all plate wells of all control plates (plates kept at 37°C). **C-D**) Mean baseline-detrended bioluminescence profiles from plate wells containing either *PER2::LUC* U2OS cells with (WT, left panel) or without *REV-ERBα* (KO, right panel) recorded at 37°C before and after transfer to 18°C for 24h (C) or 5h (D) (grey region) from biological repeat 2. 6 differently colored profiles represent cells synchronized at 6 distinct phases of the circadian period prior to the start of 18°C exposure (time zero). Black horizontal bar marks the peak "x" used in Fig. 5D. **E**) As in D but from an additional experiment in which cells were instead synchronized at 4 distinct phases of the circadian period prior to the start of 18°C exposure (4 differently colored profiles). Black horizontal bar marks the peak "x" used in F. **F**) Boxplots and individual points for times measured as in Fig. 5C for each plate well profile for both WT and KO cells kept at 37°C or transferred to 18°C for 5h from the additional experiment with four distinct profile phases prior to 18°C exposure (E and I). Points are colored according to their distinct phase. Colored lines show the change in the mean for the points from each phase. Pairwise comparisons test the significance of the difference in variance (Brown-Forsythe test (adjusted for multiple testing)) (ns = $p > 0.05$; * = $1x10^{-10} < p < 0.05$; ** = $1x10^{-20} < p < 1x10^{-10}$; *** = $p < 1x10^{-20}$). **G**) Mean baseline-detrended bioluminescence profiles from plate wells containing either *PER2::LUC* U2OS cells with (WT, left panel) or without *REV-ERBα* (KO, right panel) recorded continuously at 37°C as controls for profiles from cells transferred to 18°C for 24h (grey region). Upper two panels, biological repeat 1; lower two panels, biological repeat 2. 6 differently colored profiles for each repeat are controls for the 6 differently colored profiles from 18°C exposed plates. Red, green and blue profiles are 12h shifted versions of the pink, orange and brown profiles, respectively. Black horizontal bar marks peak "x" used in Fig. 5D (This is the second peak after the end of the grey region). **H**) As in G but control profiles for cells transferred to 18°C for 5h (grey region). Black horizontal bar marks peak "x" used in Fig. 5D (This is the second peak after the start of the grey region). **I**) As in H but control profiles for the 4 profiles from the additional experiment in E. Green and blue profiles are 6h shifted versions of the brown and purple profiles, respectively. Black horizontal bar marks peak "x" used in F (This is the second peak after the start of the grey region).

### Figure S5

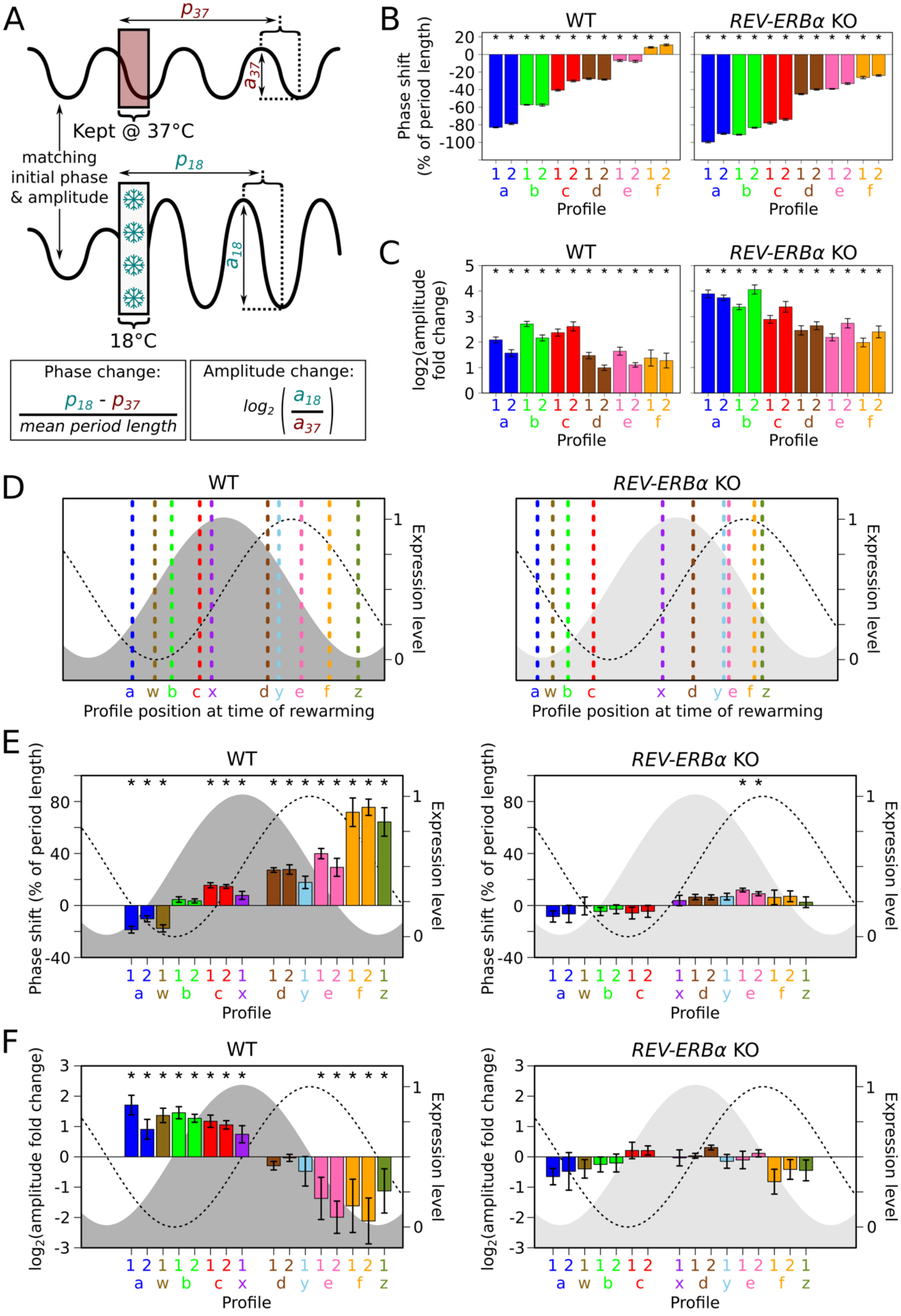
A
$p_{37}$
$a_{37}$
Kept @ 37°C
matching
initial phase
& amplitude
$p_{18}$
$a_{18}$
18°C
Phase change:
$\frac{p_{18} - p_{37}}{mean\ period\ length}$
Amplitude change:
$log_2\left(\frac{a_{18}}{a_{37}}\right)$
B
WT
REV-ERBα KO
Phase shift (% of period length)
Profile
C
WT
REV-ERBα KO
log2(amplitude fold change)
Profile
D
WT
REV-ERBα KO
Expression level
Profile position at time of rewarming
E
WT
REV-ERBα KO
Phase shift (% of period length)
Expression level
Profile
F
WT
REV-ERBα KO
$log_2$(amplitude fold change)
Expression level
Profile

**Fig. S5**

**A**) Schematic showing the calculation of the change in phase and amplitude of a profile from cells following 18°C exposure in relation to its control profile from cells kept at 37°C. **B-C**) Bar graphs showing the mean phase (B) and amplitude (C) change following 18°C exposure for 24h for each group of plate well profiles (grouped according to their distinct phase (a-f) prior to 18°C exposure) for both WT and KO cells from biological repeats 1 and 2. **D**) Schematic showing the position of the control profile (dashed line, colored according to its corresponding 5h 18°C-exposed profile) at the point of rewarming within the circadian period that is represented by the *PER2:LUC* (dashed black line) and predicted REV-ERBα expression level (grey shaded region (light grey indicates *REV-ERBα* deletion)). **E** and **F**) Bar graphs showing the mean phase (E) and amplitude (F) change following 18°C exposure for 5h for each group of plate well profiles (grouped according to their distinct phase (a-f and w-z) prior to 18°C exposure) for both WT and KO cells from biological repeats 1 and 2 using the formulas in A. Mean changes are plotted to approximately align with the position of the control profile at the point of rewarming (see D). Error bars (B-C and E-F) show the standard deviation. Asterisks mark mean phase shifts significantly greater than +/-5 % (B and E) and mean log2 amplitude fold changes significantly greater than +/-log2(1.4) (C and F) (adjusted $p < 0.05$). F is an extended version of Fig. 5E. (see Fig. 5 extended data for more details).

### Figure 5 extended data

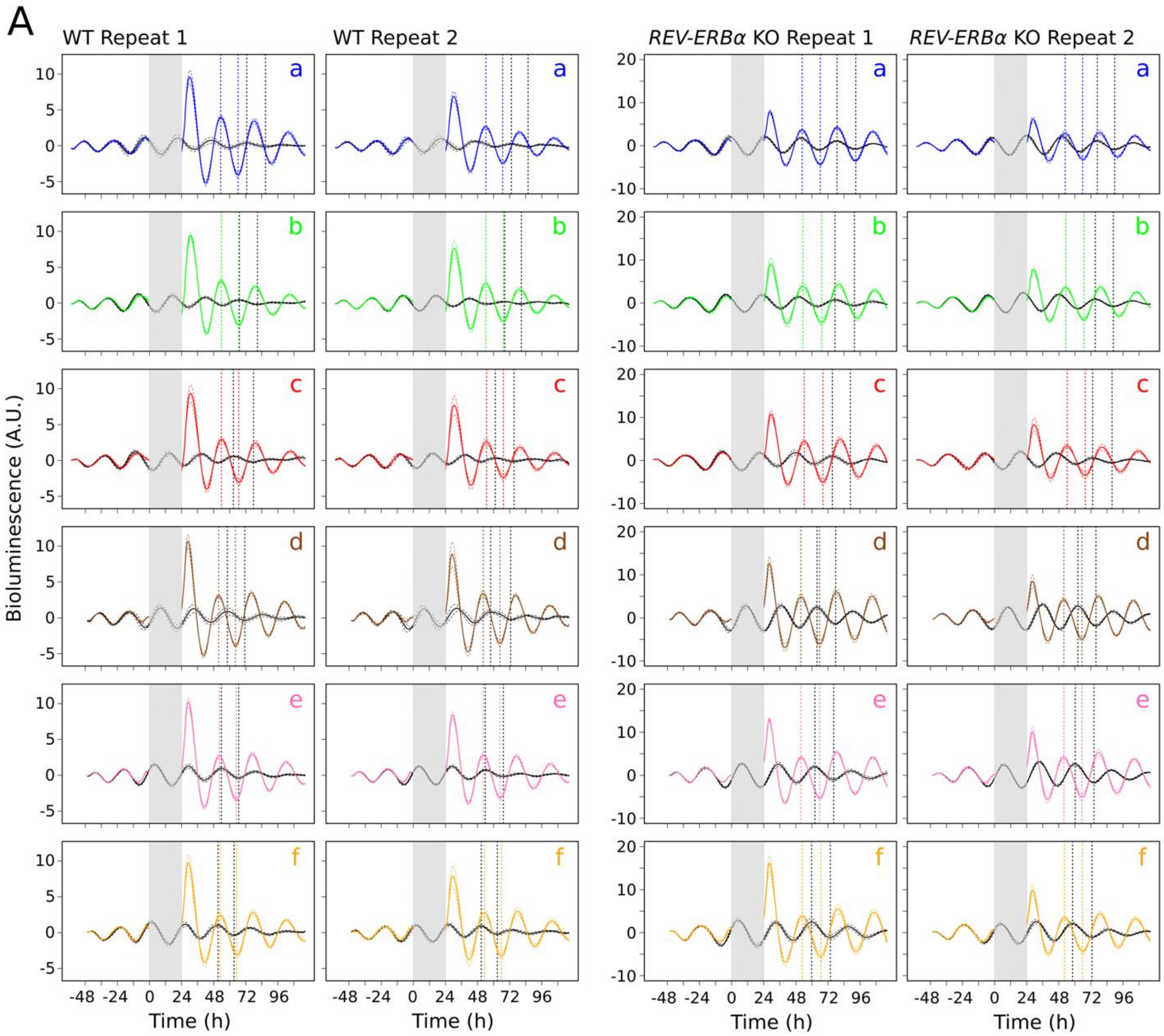

### Figure 5 extended data (continued)

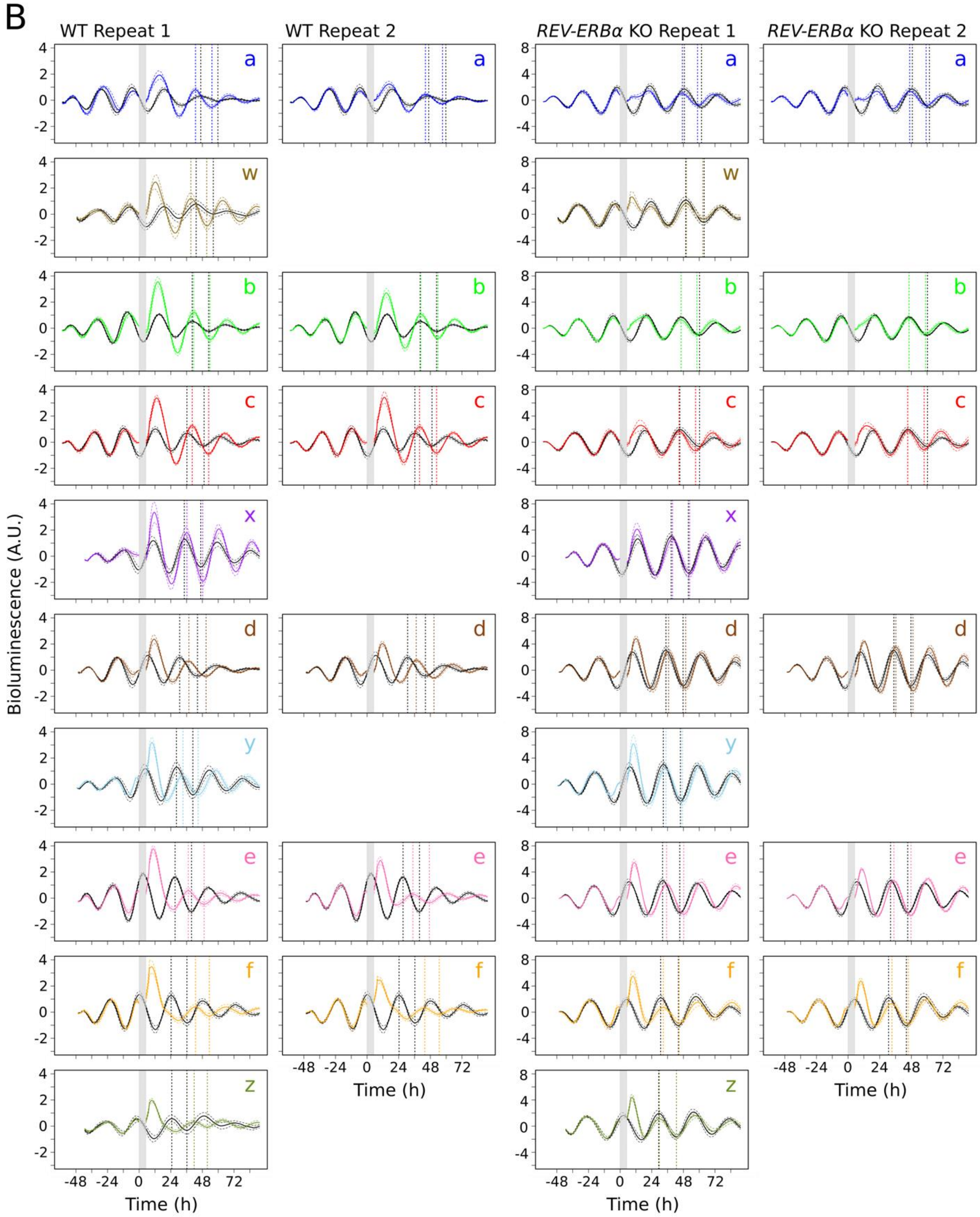

**Fig. 5 extended data**

**A-B**) Mean (solid colored profile) and standard deviation (dashed colored profile) of baseline-detrended bioluminescence profiles from plate wells containing either *PER2::LUC* U2OS cells with (WT, leftmost 2 panels) or without *REV-ERBα* (KO, rightmost 2 panels) recorded at 37°C before and after transfer to 18°C for 24h (A) or 5h (B) (grey region) compared to control profiles kept and recorded continuously at 37°C (black solid (mean) and dashed (standard deviation) profiles) from two biological repeats. 6 differently colored profiles (a-f) represent cells synchronized at 6 distinct phases of the circadian period prior to the start of 18°C exposure (time zero) (A and B). 4 differently colored profiles (w-z) from an additional experiment (Fig. S4E,I) represent cells synchronized at 4 distinct phases of the circadian period prior to the start of 18°C exposure (B only). Dotted vertical lines mark the peaks and troughs of the control (black) and 18°C-exposed (colored) profiles used to calculate phase and amplitude changes in Fig. S5B,E and Fig. 5E, S5C,F, respectively, using the formulas in Fig. S5A.

References:

Kuleshov, M. V, Jones, M.R., Rouillard, A.D., Fernandez, N.F., Duan, Q., Wang, Z., Koplev, S., Jenkins, S.L., Jagodnik, K.M., Lachmann, A., et al. (2016). Enrichr: a comprehensive gene set enrichment analysis web server 2016 update. Nucleic Acids Res. *44*, W90–W97.

Love, M.I., Huber, W., and Anders, S. (2014). Moderated estimation of fold change and dispersion for RNA-seq data with DESeq2. Genome Biol. *15*, 550.

Lugowski, A., Nicholson, B., and Rissland, O.S. (2018). Determining mRNA half-lives on a transcriptome-wide scale. Methods *137*, 90–98.
